# Essential functions of mosquito ecdysone importers in development and reproduction

**DOI:** 10.1101/2021.09.10.459809

**Authors:** Lewis V. Hun, Naoki Okamoto, Eisuke Imura, Roilea Maxson, Riyan Bittar, Naoki Yamanaka

**Affiliations:** Department of Entomology, Institute for Integrative Genome Biology, University of California, Riverside, 900 University Ave., Riverside, CA 92521, USA; Life Science Center for Survival Dynamics, Tsukuba Advanced Research Alliance, University of Tsukuba, Tsukuba, Ibaraki 305-8577, Japan

## Abstract

The primary insect steroid hormone ecdysone requires a membrane transporter to enter its target cells. Although an organic anion-transporting polypeptide (OATP) named Ecdysone Importer (EcI) serves this role in the fruit fly *Drosophila melanogaster* and most likely in other arthropod species, this highly conserved transporter is apparently missing in mosquitoes. Here we report three additional OATPs that facilitate cellular incorporation of ecdysone in *Drosophila* and the yellow fever mosquito *Aedes aegypti*. These additional ecdysone importers (EcI-2, 3, and 4) are dispensable for development and reproduction in *Drosophila*, consistent with the predominant role of EcI. In contrast, in *Aedes*, EcI-2 is indispensable for ecdysone-mediated development, whereas EcI-4 is critical for vitellogenesis induced by ecdysone in adult females. Altogether, our results indicate unique and essential functions of these additional ecdysone importers in mosquito development and reproduction, making them attractive molecular targets for species- and stage-specific control of ecdysone signaling in mosquitoes.

## INTRODUCTION

Ecdysone and other ecdysteroids are a group of steroid hormones that control various aspects of insect development and reproduction (1). Once released into the hemolymph, ecdysone (more specifically, its active form 20-hydroxyecdysone or 20E and related ecdysteroids) enters its target cells to bind to a nuclear receptor named the ecdysone receptor (EcR), which forms a heterodimer with another nuclear receptor Ultraspiracle and induces gene expression (2–5). Although it has long been assumed that ecdysone can pass through the cell membrane through simple diffusion, we recently demonstrated that an organic anion-transporting polypeptide (OATP), which we named Ecdysone Importer (EcI), is required for cellular uptake of ecdysone in the fruit fly *Drosophila melanogaster* (6). *EcI* orthologs are found among a wide variety of insects (6), and it is likely that its critical function as a mediator of ecdysone signaling is highly conserved in other insect species (7).

Mosquitoes are the deadliest disease vectors for humans. The yellow fever mosquito *Aedes aegypti* is the primary vector for arboviruses including Zika, yellow fever, chikungunya and dengue viruses, which are of global health concern due to their rapid increases in the geographical distribution (8, 9). Critical functions of ecdysone signaling in *Aedes* development and reproduction are extensively investigated (10–13), making it an important molecular target for control agents against this deadly virus vector. Surprisingly, however, an *EcI* ortholog cannot be found in well-annotated mosquito genomes (14–19), suggesting the existence of additional membrane transporter(s) for ecdysone.

In this study, we identified additional OATPs that facilitate cellular uptake of ecdysone in *Drosophila* and *Aedes*. These additional ecdysone importer-encoding genes, *EcI-2*, *3*, and *4*, are dispensable for normal development and reproduction in *Drosophila*, confirming the predominant role of *EcI* in flies. In contrast, *CRISPR/Cas9*-mediated mutagenesis and RNA interference (RNAi)-mediated knockdown experiments in *Aedes* suggest that *EcI-2* is critical for ecdysone-mediated developmental progression, while *EcI-4* is most important for vitellogenesis induced by ecdysone in adult females. Collectively, our results indicate unique functions of these additional ecdysone importers in mosquitoes, making them attractive targets for species- and stage-specific control of mosquito ecdysone signaling.

## RESULTS

### Identification of additional ecdysone importers in *Drosophila* and *Aedes*

Our detailed phylogenetic analysis of dipteran OATPs confirmed absence of *EcI* orthologs in mosquitoes (Figure 1A). To identify additional ecdysone importers, we used an ecdysteroid-inducible gene expression system in HEK293 cells (20–22) to test if the other OATPs in *Drosophila* and *Aedes* can facilitate ecdysone signaling in a heterologous system. This led to identification of three additional ecdysone importers in each species: Oatp33Ea, Oatp58Dc, and Oatp58Db in *Drosophila*, and AAEL007691, AAEL010921, and AAEL010917 in *Aedes* (Figure 1B). Based on phylogenetic similarities, these additional ecdysone importers in each species were named EcI-2, 3, and 4, respectively.

**Figure 1.**
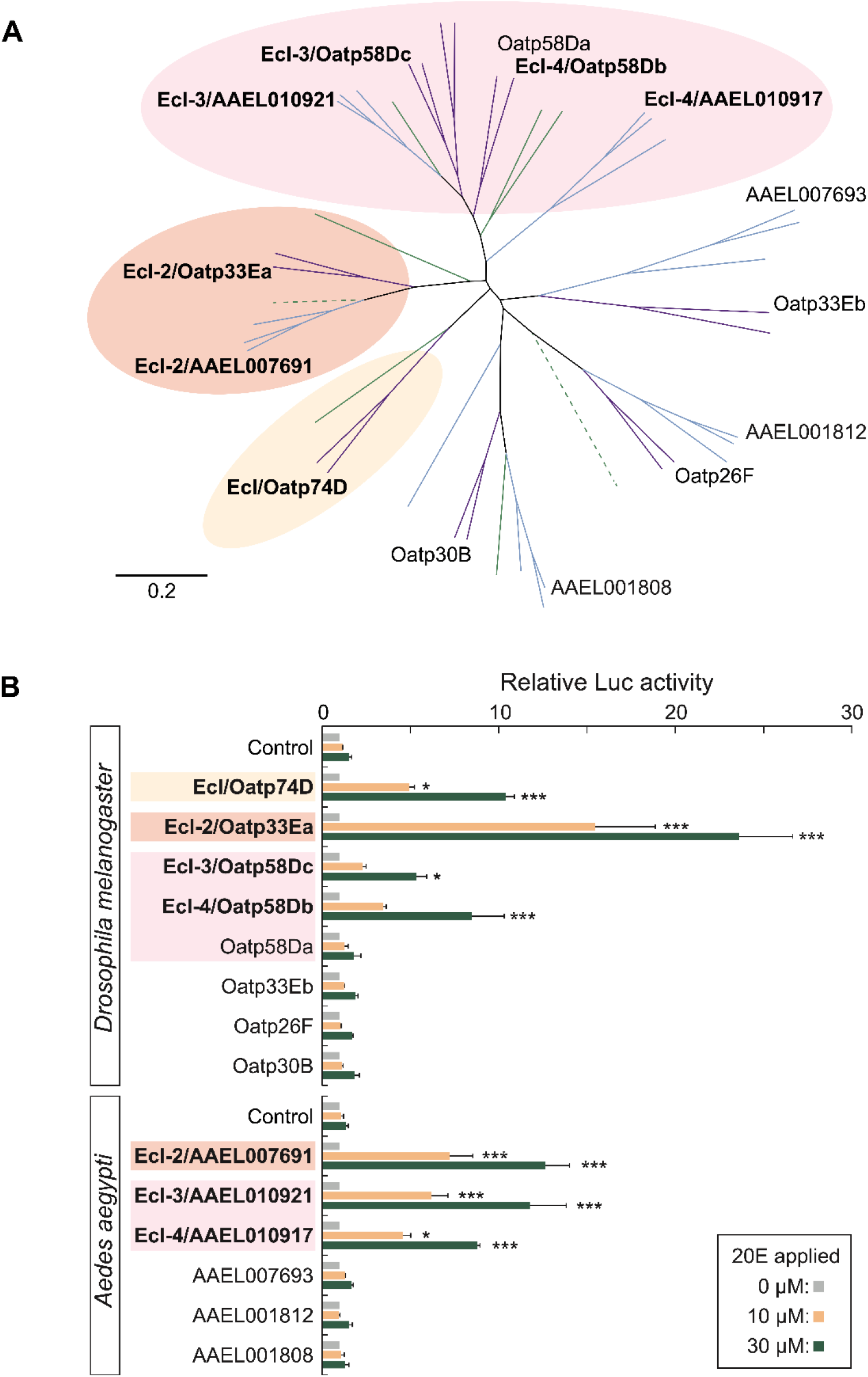
Identification of additional ecdysone importers in *Drosophila* and *Aedes*. (A) Neighbor-joining unrooted phylogenetic tree of full length OATP proteins from representative dipteran insect species. Purple, flies (*Drosophila melanogaster* and *Musca domestica*); green, sand flies (*Phlebotomus papatasi*); blue, mosquitoes (*Aedes aegypti*, *Anopheles gambiae*, and *Culex quinquefasciatus*). Dotted lines indicate pseudogenes. *Drosophila* and *Aedes* OATPs are labeled. Protein names and GenBank accession numbers are listed in Table S5. Scale bar indicates an evolutionary distance of 0.2 amino acid substitutions per position. (B) Luciferase (Luc) reporter activities in response to 10 μM or 30 μM 20E in HEK293 cells expressing *Drosophila* or *Aedes* OATPs. The cells were transfected with modified *EcR* (*VgEcR*) and *RXR*, along with each OATP-containing vector and luciferase reporter plasmids. Values are relative to the basal level (0 M 20E). All values are the means ± SEM (n = 2-4). *p < 0.05, ***p < 0.001 from one-way ANOVA followed by Dunnett’s multiple comparison test as compared to the response of the control cells to the same concentration of 20E.

Temporal expression of *EcI-2*, *3*, and *4* fluctuates during development in flies and mosquitoes (Figure S1A and B). With regard to tissue-specific expression, we previously showed that *EcI-2* is highly expressed in the gut, whereas *EcI-3* and *EcI-4* are predominantly expressed in the Malpighian tubules in *Drosophila* larvae (6). Consistent with this, we observed high expression of *EcI-2* in the gut and *EcI-3* and *EcI-4* in the Malpighian tubules, respectively, in *Aedes* larvae (Figure S1C). Importantly, albeit its strong expression in the gut, *Aedes EcI-2* is expressed at higher levels in the head and carcass as compared to the other ecdysone importers, suggesting its potential ubiquitous function during the larval stage (Figure S1C).

### Unique requirement of additional ecdysone importers during mosquito embryogenesis

We next conducted *CRISPR/Cas9*-mediated mutagenesis of *EcI-2*, *3*, and *4* in *Drosophila* and *Aedes*, aiming to investigate their functions during the life cycle. In *Drosophila*, neither individual mutants nor combination mutants showed any discernible developmental or reproductive defects (Figure S2 and S3, Table S1 and S2), consistent with the predominant role of *EcI* as an ecdysone importer in flies (6). In contrast, when single guide RNA (sgRNA) for each gene (Figure S4) was injected with Cas9 protein into *Aedes* eggs, all ecdysone importer mutants showed significant embryonic lethality as compared to controls (Figure 2A). Importantly, only *EcI-2* mutants showed levels of embryonic lethality comparable to *EcR* knockout animals (Figure 2A). Some of the *EcI-2* mutants also showed hatching defects similar to *EcR* mutants (Figure 2B), suggesting a critical function of EcI-2 in ecdysone signaling during mosquito embryogenesis.

**Figure 2.**
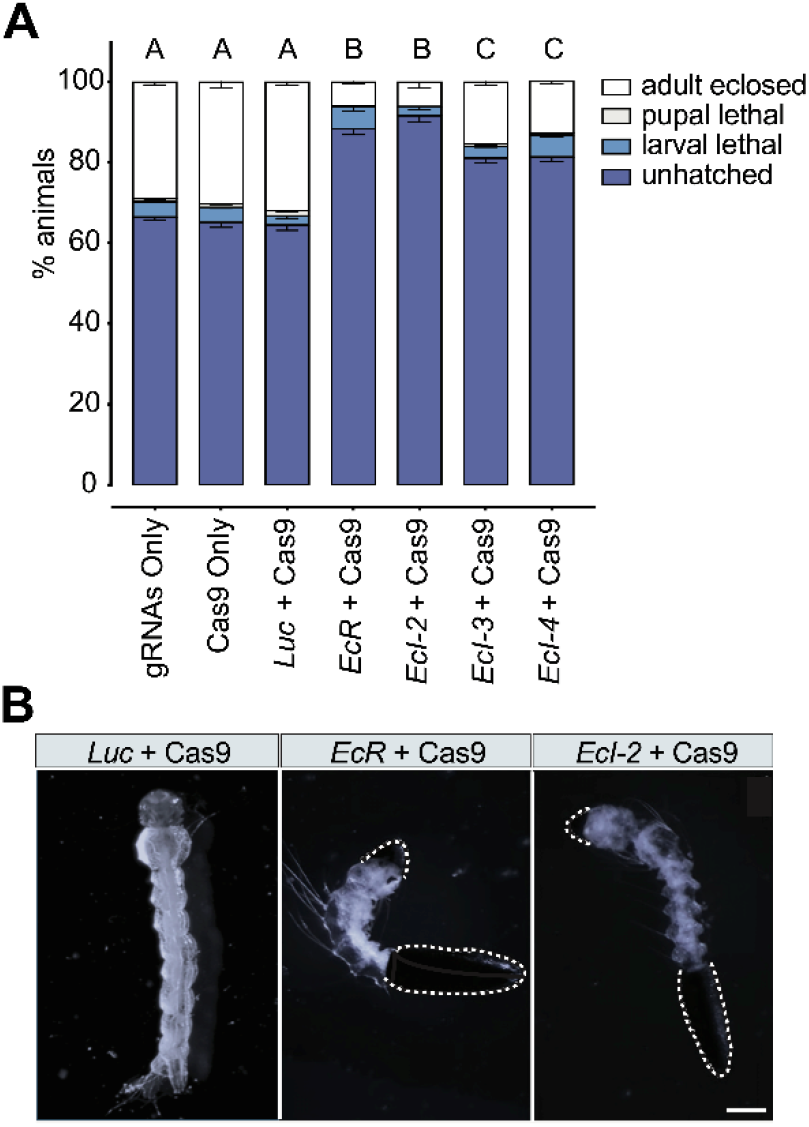
*Aedes* ecdysone importers are required for embryogenesis. (A) Lethal stages of *Aedes* embryos injected with Cas9 protein and sgRNAs against control (*Luc*), *EcR*, or ecdysone importer genes. Total 450 eggs were injected in three independent batches for each treatment, and their lethal stages were monitored thereafter. All values are the means ± SEM. Same letters on top of the bars indicate statistically insignificant differences in embryonic lethality based on ANOVA with Tukey’s HSD (honestly significant difference) test. (B) Hatching defects observed in *EcR* and *EcI-2* mutants. As compared to control (*Luc* + Cas9) that hatched normally, *EcR* or *EcI-2* mutagenesis caused hatching defects in some larvae, where the eggshell remained attached (dotted white lines). Scale bar, 0.5 mm.

### *EcI-2* is required for mosquito larval development

In order to circumvent embryonic lethality in ecdysone importer mutants, we undertook an RNAi approach to further investigate their potential involvement in ecdysone signaling at later developmental stages. Newly hatched *Aedes* larvae were soaked in double strand RNA (dsRNA) solutions to suppress expression of each ecdysone importer or *EcR*, which resulted in ~50-60% reduction of mRNA levels for each target gene (Figure S6). Periodic monitoring of their developmental stages and lethality revealed that *EcR* knockdown induced high (80-90%) lethality beginning 48 hours after hatching (hAH), consistent with the critical function of ecdysone signaling in molting induction during larval stages (Figure 3A). Likewise, knockdown of *EcI-2*, but not *EcI-3* or *EcI-4*, induced 70-80% lethality during larval development. Importantly, after 68 hAH, many surviving *EcR* RNAi and *EcI-2* RNAi larvae remained as younger instars compared to control animals (Figure 3A), and their size increase was clearly inhibited (Figure 3B). This developmental arrest phenotype in *EcR* RNAi and *EcI-2* RNAi animals was further confirmed by detailed measurement of head size and siphon length at two different time points during larval development (Figure S5). Furthermore, expression of an ecdysone-inducible gene, *E74B*, was suppressed by ~70% in *EcI-2* RNAi animals, as compared to milder (~40%) reduction in *EcI-3* or *EcI-4* RNAi animals (Figure S7). Importantly, this strong inhibition of *E74B* expression in *EcI-2* RNAi animals was comparable to RNAi knockdown of *EcR* or *shade*, the gene encoding a cytochrome P450 enzyme required for 20E production (23, 24). Overall, these results further suggest involvement of EcI-2 in ecdysone signaling during mosquito development.

**Figure 3.**
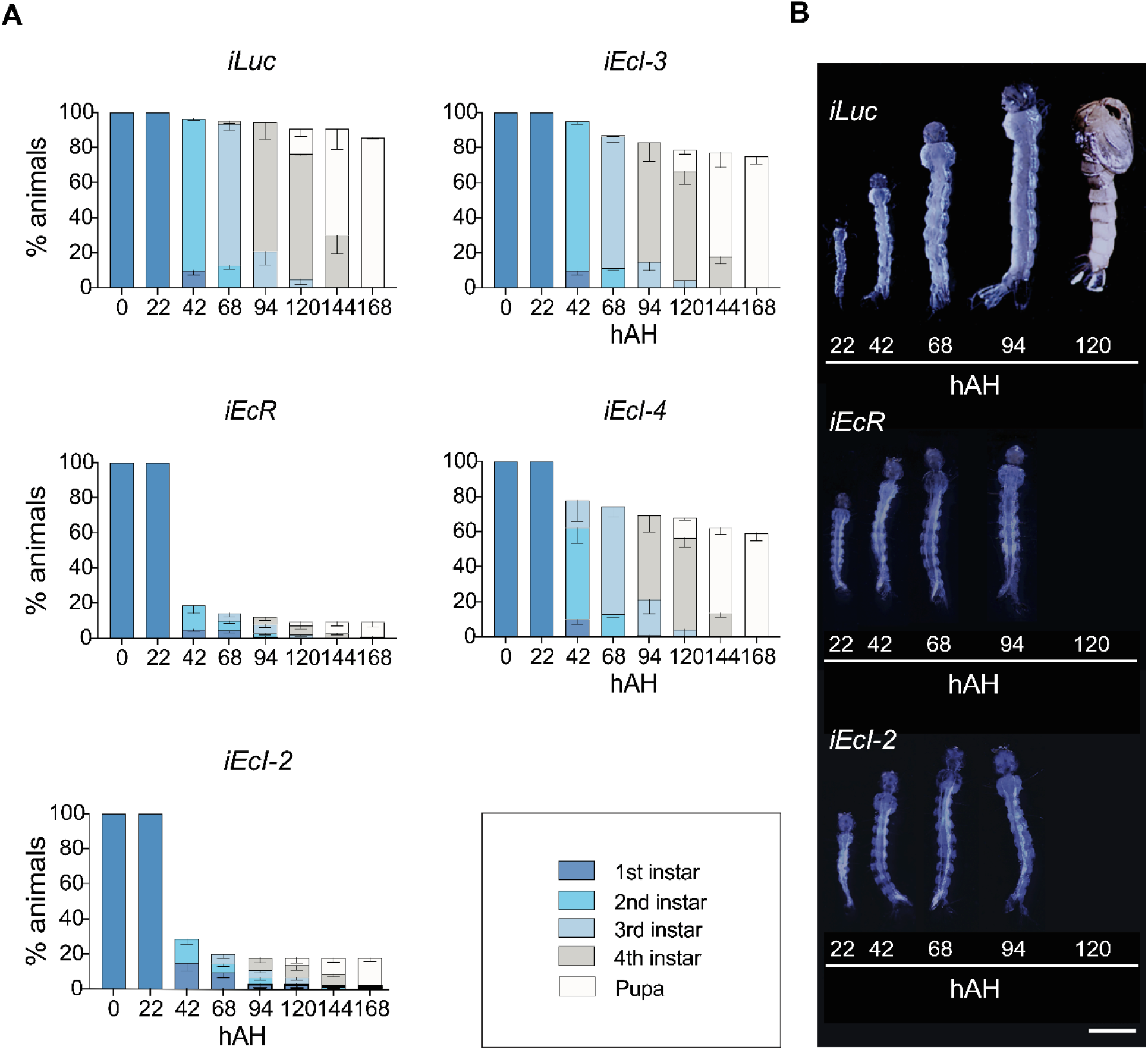
*Aedes EcI-2* is required for larval developmental transitions. (A) Developmental progression and survival rate (%) of *Luc* RNAi (*iLuc*; control), *EcR* RNAi (*iEcR*), and *EcI-2*, *3*, and *4* RNAi (*iEcI-2*, *iEcI-3*, and *iEcI-4*) animals. Color bars indicate developmental stages determined by stage-specific morphologic features such as the head capsule size and siphon length (28, 29). All values are the means ± SEM from 7 independent experiments with 20 individuals in each replicate. (B) Developmental progression of animals treated with dsRNA targeting *Luc* (*iLuc*; control), *EcR* (*iEcR*), or *EcI-2* (*iEcI-2*). Representative images of animals at various time points were combined into single panels. Scale bar, 1 mm.

We next attempted to rescue the larval arrest phenotype by administration of two different EcR agonists: the endogenous ecdysteroid 20E and a non-steroidal insecticide chromafenozide (CF) (Figure 4). As we have previously shown that CF enters cells independently of EcI (6, 25), we reasoned that CF can activate EcR even in the absence of ecdysone importers, thereby rescuing the developmental defect caused by *EcI-2* knockdown. Indeed, CF but not 20E significantly rescued the larval arrest phenotype in *EcI-2* RNAi animals, whereas these agonists only partially rescued the growth defect, if at all, caused by *EcR* knockdown (Figure 4). Importantly, both CF and 20E significantly rescued developmental arrest caused by knockdown of *shade*, confirming the potency of these compounds as EcR agonists *in vivo*. Collectively, these results indicate that EcR remains functional in the absence of EcI-2, and that EcI-2 is required for 20E to access EcR and induce larval development in mosquitoes.

**Figure 4.**
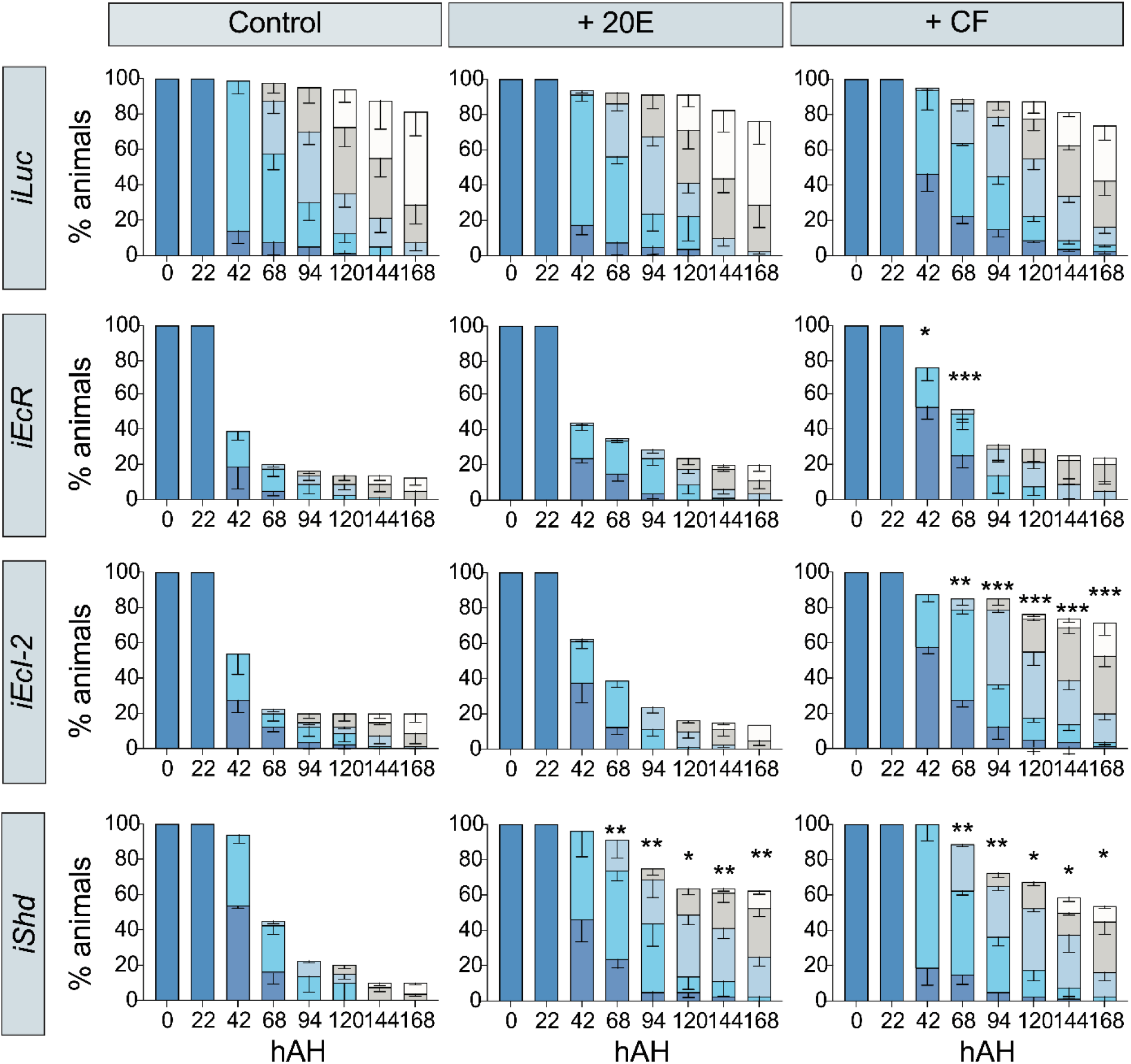
A non-steroidal ecdysone agonist can rescue developmental arrest caused by *EcI-2* knockdown in *Aedes*. Developmental progression and survival rate (%) of *Luc* RNAi (*iLuc*; control), *EcR* RNAi (*iEcR*), *EcI-2* RNAi (*iEcI-2*), and *shade* RNAi (*iShd*) animals treated with 20E or chromafenozide (CF). Color bars indicate developmental stages as shown in Figure 3A. All values are the means ± SEM from 4 independent experiments with 20 individuals in each replicate. *p < 0.05, **p < 0.01, ***p < 0.001 from one-way ANOVA followed by Bonferroni’s multiple comparison test as compared to the survival rate of control (no agonist treatment) of the same RNAi animal at the same time point.

### *EcI-4* is required for mosquito vitellogenesis

In adult female mosquitoes, vitellogenesis after the blood meal is primarily controlled by 20E. 20E upregulates expression of ecdysone-inducible genes, whose products in turn promote expression of yolk protein precursors such as vitellogenin (Vg) in the fat body (10, 11). When expression levels of ecdysone importers were periodically monitored after the blood meal, we observed significant fluctuation of their expression in all tissues observed, potentially reflecting dynamic changes in ecdysone signaling after blood feeding (Figure S8). Importantly, predominant expression of *EcI-3* and *EcI-4* was observed in the abdomen, where the fat body is located (Figure S8).

We next injected dsRNA targeting each ecdysone importer or *EcR* into newly eclosed females, which were blood-fed 3 days after injection. This resulted in ~60-75% reduction of mRNA levels for each target gene (Figure S9). Vitellogenesis was clearly reduced in *EcR* RNAi animals as indicated by ovary length, follicle size, and yolk length (Figure 5A-D), consistent with the strong reduction of *Vg* expression in the fat body (Figure S9E). Importantly, *Vg* expression was also strongly reduced in *EcI-4* RNAi animals, as compared to milder reduction caused by *EcI-2* or *EcI-3* knockdown (Figure S9E). Consistent with this, knockdown of *EcI-4*, but not *EcI-2* or *EcI-3*, caused similar reduction in vitellogenesis comparable to that caused by *EcR* knockdown (Figure 5A-D). As a result, both *EcR* RNAi and *EcI-4* RNAi caused significant reduction in fecundity (Figure 5E). Taken together, our results indicate the predominant role of EcI-4 in ecdysone-mediated reproduction in adult female mosquitoes.

**Figure 5.**
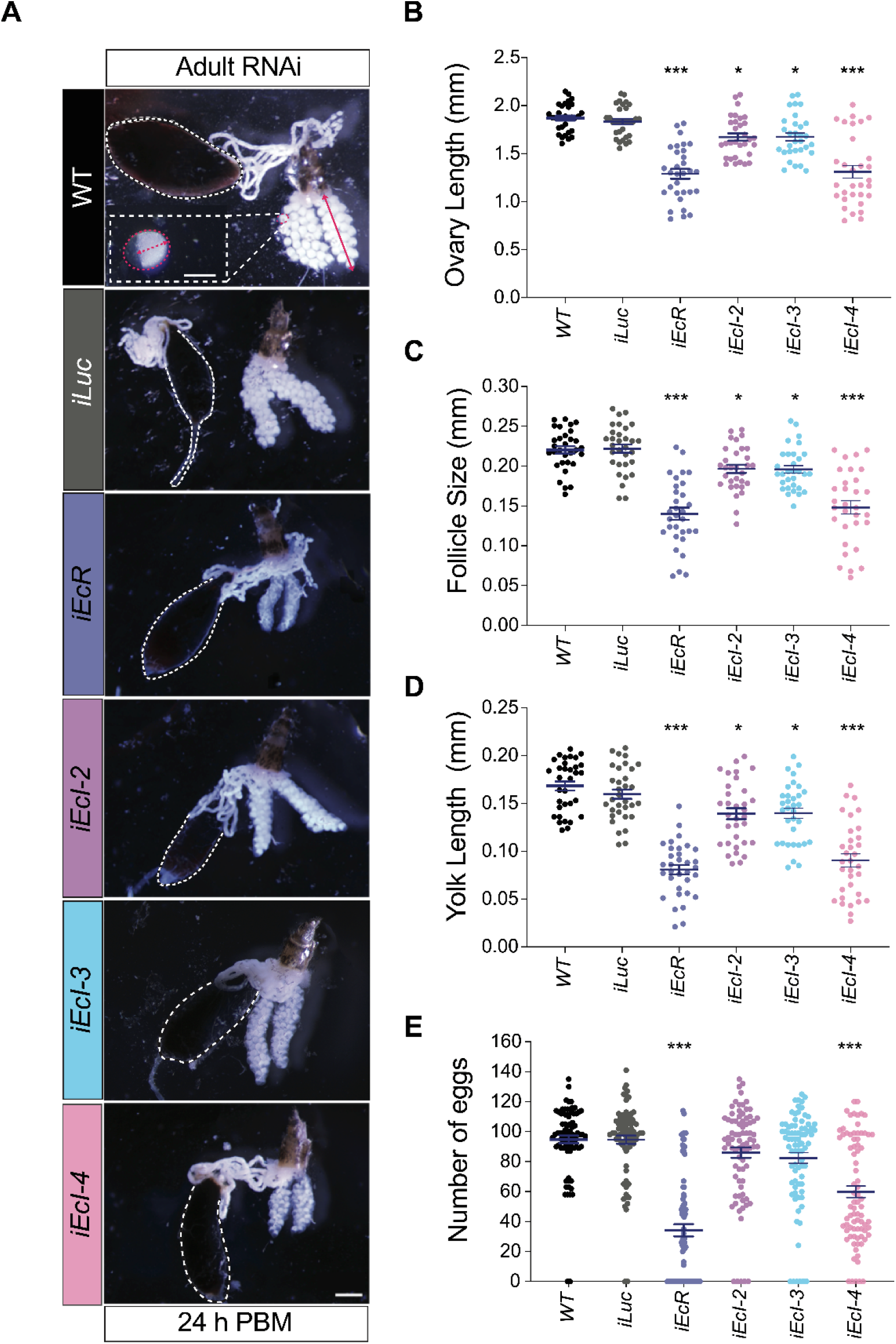
*EcI-4* is required for vitellogenesis in *Aedes* adult females. (A) Representative images of ovaries from wild type (*WT*), *Luc* RNAi (*iLuc*; control), *EcR* RNAi (*iEcR*), and *EcI-2*, *3*, and *4* RNAi (*iEcI-2*, *iEcI-3*, and *iEcI-4*) females 24 hours post blood meal (PBM). The gut filled with the blood meal is surrounded by dashed white lines. In the top panel, ovary length, follicle size, and yolk size are indicated by solid red arrow, dashed red line (longitudinal axis), and dashed red arrow, respectively. Scale bars, 150 μm for follicle and 0.5 mm for ovary. (B-D) Ovary length (B), follicle size (C), and yolk length (D) of wild type (*WT*), *Luc* RNAi (*iLuc*; control), *EcR* RNAi (*iEcR*), and *EcI-2*, *3*, and *4* RNAi (*iEcI-2*, *iEcI-3*, and *iEcI-4*) females. All values are the means ± SEM from a minimum of 3 independent experiments with 10 individuals in each replicate. *p < 0.05, ***p < 0.001 from one-way ANOVA followed by Dunnett’s multiple comparison test as compared to control. (E) Number of deposited eggs per each individual of wild type (*WT*), *Luc* RNAi (*iLuc*; control), *EcR* RNAi (*iEcR*), and *EcI-2*, *3*, and *4* RNAi (*iEcI-2*, *iEcI-3*, and *iEcI-4*) females. All values are the means ± SEM from a minimum of 3 independent experiments with 10 individuals in each replicate. ***p < 0.001 from one-way ANOVA followed by Dunnett’s multiple comparison test as compared to control.

## DISCUSSION

Recent studies in *Drosophila* have shown that ecdysone, the primary steroid hormone in insects, requires a membrane transporter EcI to enter its target cells and exert its genomic effects through EcR (6, 25). Although this critical function of EcI in ecdysone signaling seems to be highly conserved among other insect species (7), we noticed that *EcI* orthologs are unexpectedly missing in mosquitoes. In the present study, we characterized additional ecdysone importers, EcI-2, 3, and 4, in both *Drosophila* and *Aedes*. These additional ecdysone importers are dispensable for development and reproduction in *Drosophila*, likely reflecting the predominant role of EcI as an ecdysone importer. In contrast, in *Aedes* mosquitoes, EcI-2 is required for ecdysone-mediated developmental progression, whereas EcI-4 seems to be most critical for vitellogenesis in adult females.

The evolutionary reason why *EcI* was lost in mosquito species is currently unknown. Our results nonetheless suggest that, in the absence of *EcI*, alternative ecdysone importers assume paramount importance in development and reproduction. This makes these additional ecdysone importers potential targets for development of mosquito-specific insect growth regulators and pesticides. The involvement of distinct ecdysone importers in development and reproduction may even make it possible to develop stage-specific regulators of ecdysone signaling in mosquitoes. The effort to identify efficient blockers of ecdysone importers is currently underway.

What are the functions of the additional ecdysone importers in other insects? Although *EcI-2*, *3*, and *4* are dispensable for *Drosophila* development and reproduction, it is conceivable that they have either unique or redundant functions that are not essential for overall viability and fertility under normal conditions. In this regard, it would be important to examine potential functions of these additional ecdysone importers in *Drosophila* and other insects more closely under different conditions, particularly in tissues where these transporters are highly expressed (6). Considering pleiotropic functions of ecdysone in insect physiology (1), it would also be interesting to investigate whether these additional ecdysone importers are essential for ecdysone functions that are not directly related to growth and reproduction, such as stress responses. Some tools developed in the current study, such as *EcI-2*, *3*, and *4* mutant flies, are expected to facilitate such future studies.

In conclusion, our results indicate unique functions of newly characterized ecdysone importers in development and reproduction in mosquitoes, which may pave the way for better control of these deadly human disease vectors. Elucidating as yet unknown functions of related ecdysone importers in other insect species is expected to deepen our understanding of how pleiotropic functions of ecdysone are regulated in different tissues under different conditions.

## MATERIALS AND METHODS

### Flies

All flies (*Drosophila melanogaster*) were raised at 25°C on standard fly food (6) under 12-hour light/dark cycle. *w*^*1118*^ was used as a control strain. *EcI* mutant alleles (*EcI*^*1*^ and *EcI*^*2*^) were generated previously (6); all the other ecdysone importer mutant alleles (*EcI-2*^*1*^, *EcI-3*^*1*^, *EcI-4*^*1*^, and *EcI-3-4*^*1*^) were generated using CRISPR/Cas9-mediated mutagenesis as described below. Deficiency alleles over ecdysone importer genes (Df(2L)Exel6033, #7516; Df(2R)Exel7171, #7902) were obtained from the Bloomington *Drosophila* Stock Center (BDSC).

### Mosquitoes

The Rockefeller strain of *Aedes aegypti* was used in this study. Both male and female adults were maintained on 10% sucrose and reared at 27°C and 80% relative humidity under 16-hour light/8-hour dark cycle. Larvae in water were fed on a diet containing dry dog food (Blue Buffalo), fish flakes (Tetramin), and liver powder (Now Foods) (10:10:1 wt. ratio). All mosquito dissection was performed in 1X phosphate-buffered saline (PBS) solution (Fisher BioReagents) at room temperature. Using an artificial glass feeder, all adult female mosquitoes were allowed to feed on bovine blood purchased from Hemostat. Only fully engorged female mosquitoes were used.

### Phylogenetic tree analysis

The unrooted neighbor-joining tree (Figure 1A) was generated using ClustalW (DNA Data Bank of Japan). Entire amino acid sequences of all OATPs in the following dipteran species were aligned: the fruit fly *Drosophila melanogast*er, the housefly *Musca domestica*, the sand fly *Phlebotomus papatasi*, the yellow fever mosquito *Aedes aegypti*, the African malaria mosquito *Anopheles gambiae*, and the southern house mosquito *Culex quinquefasciatus*. Protein names and GenBank accession numbers are listed in Table S3.

### Cloning of OATP-encoding genes

For *Drosophila* OATP-encoding genes, full length cDNA clones from *Drosophila* Genomics Resource Center (*EcI-2/Oatp33Ea*, LD36578; *EcI-3/Oatp58Dc*, LP09443; *Oatp58Da*, IP17768; *Oatp33Eb*, RE09129; *Oatp26F*, RE32029; *Oatp30B*, RE26532) were subcloned into *pcDNA3.1* vector. The coding sequence of *EcI-4/Oatp58Db* was PCR-amplified from whole-body *Drosophila* cDNA using Phusion Plus DNA Polymerase (Thermo Fisher Scientific) with the primers listed in Table S4. The PCR product was cloned into the *pcDNA3.1* vector and sequenced.

For *Aedes* OATP-encoding genes, total RNA was collected from 4th instar larvae using TRIzol reagent (Invitrogen) according to the manufacturer’s instructions. cDNA was generated from purified total RNA using PrimeScript RT Master Mix (Takara Bio). PCR amplification of the sequence corresponding to *EcI-2/AAEL007691*, *EcI-3/AAEL010921*, *EcI-4/AAEL010917*, *AAEL001812*, *AAEL001808*, *AAEL007693* was performed using Phusion Plus DNA Polymerase with the primers listed in Table S5. The PCR products were cloned into the *pcDNA3.1* vector and sequenced.

### Transfection and luciferase reporter assay in HEK293 cells

HEK cells (obtained from Michael E. Adams) at a density of 4 × 10^5^ cells/ml were seeded in 100 μl/well of Opti-MEM reduced serum media (Thermo Fisher Scientific) containing 5% FBS and 1% MEM non-essential amino acids (NEAA) solution (Thermo Fisher Scientific) in a 96-well clear flat bottom microplate (Corning). Transfection of HEK293 cells was performed using Attractene transfection reagent (Qiagen) by the fast-forward transfection approach following the manufacturer’s instructions. 50 μl/well of transfection cocktail containing Opti-MEM reduced serum media, Attractene, and DNA plasmids was added to each well, bringing the final volume to 150 μl/well. 0.1 μg/well of *pcDNA3.1* empty vector (control) or *pcDNA3.1* vector containing full length cDNA of *Drosophila* or *Aedes* OATP-encoding genes was transfected, along with 60 ng/well of *pERV3* receptor plasmid (Agilent Technologies) containing a modified ecdysone receptor (VgEcR) and RXR, 36 ng/well of *pEGSH-LUC* luciferase reporter plasmid (Agilent Technologies), and 0.9 ng/well of *pRL-CMV* Renilla luciferase reporter plasmid (Promega) as a reference. After 24 hours of incubation at 37°C and 5% CO_2_, transfection medium was removed and replaced with 150 μl/well of DMEM with 4.5 mg/ml glucose and sodium pyruvate without L-glutamine and phenol red (w-G-SP, wo-G-PR) (Thermo Fisher Scientific) containing 10% FBS, 1% PSS, and 1% MEM NEAA solution. After 48 hours of incubation at 37°C and 5% CO_2_, medium was removed and replaced with 150 μl/well of DMEM (w-G-SP, wo-G-PR) containing 1% PSS, 1% MEM NEAA solution, and 20E at indicated concentrations. After 24 hours of incubation at 37°C and 5% CO_2_, 75 μl of media was removed (total 75 μl remaining per well) and 75 μl/well of Dual-Glo Solution was added (total 150 μl/well). After 20 min of incubation at RT in the dark, 120 μl/well of cell lysates were transferred to 96-well solid white flat bottom polystyrene TC-treated plates. The firefly luciferase activity and co-transfected Renilla luciferase activity were measured subsequently using the Dual-Luciferase Reporter Assay System in accordance with the manufacturer’s instructions and analyzed with GloMax-Multi + Microplate Multimode Reader with Instinct. For each condition, cells were treated independently in 3 wells of a 96-well plate in each experiment, and the same experiment was conducted multiple times on different days.

### Total RNA extraction and quantitative reverse transcription (qRT)-PCR

Animals or dissected tissues were collected in 1.5 ml tubes and immediately flash-frozen in liquid nitrogen. Total RNA from animals or tissues was extracted using TRIzol reagent (Invitrogen) according to the manufacturer’s instructions. Extracted RNA was further purified by RNeasy mini kit (Qiagen) following the manufacturer’s instructions, combined with treatment with RNase-Free DNase Set (Qiagen). cDNA was generated from purified total RNA using PrimeScript RT Master Mix (Takara Bio). qRT-PCR was performed on the CFX connect real-time PCR detection system (Bio-Rad) using SYBR Premix Ex Taq II (Tli RNaseH Plus) (Takara Bio). For absolute quantification of mRNAs, serial dilutions of pGEM-T (Promega) plasmids containing coding sequences of the target genes or internal control genes (*rp49* for *Drosophila melanogaster* and *AeRpL32* (*AAEL003396*) for *Aedes aegypti*) were used as standards. After the molar amounts were calculated, transcript levels of the target mRNA were normalized to internal control gene levels in the same samples. Three separate samples were collected for each experiment and duplicate measurements were conducted. The primers used are listed in Table S4 and S5.

### Mutagenesis in *Drosophila*

*EcI-2*, *EcI-3*, and *EcI-4* mutant alleles were generated using the CRISPR/Cas9 system. Pairs of gRNA target sequences (20 bp: T1 and T2) were designed near the transcription start and stop sites of each target gene using NIG-FLY Cas9 Target finder (NIG) (Figure S2). T1 from *EcI-3* and T2 from *EcI-4* were used for making the *EcI-3-4* double mutant. Forward and reverse 24-bp oligonucleotides with 20-bp target sequences (Table S4) were annealed to generate a double-strand DNA with 4-bp overhangs on both ends and inserted into *BbsI*-digested *pBFv-U6.2* or *pBFv-U6.2B* vector provided by the NIG (26). To construct double-gRNA vectors, the first gRNA (T1) was cloned into *pBFv-U6.2* (named *pBFv-U6.2-T1*), whereas the second gRNA (T2) was cloned into *pBFv-U6.2B* (named *pBFv-U6.2B-T2*). A fragment containing the U6 promoter and the first gRNA was cut out from *pBFv-U6.2-T1* and ligated into *pBFv-U6.2B-T2* (named *pBFv-U6.2B-T1-T2*). These double-gRNA vectors (*pBFv-U6.2B-T1-T2*) were independently injected into embryos of *yw;; nos-cas9 (III-attP2)/TM6B* flies (BestGene Inc).

For each target gene, surviving G0 males were divided into 10 groups and crossed *en masse* to *Sp/CyO-GFP* (obtained from Takashi Nishimura) virgin female flies. From the progeny of each of these ten crosses, 5 single males were isolated and crossed independently to *Sp/CyO-GFP* virgin female flies to establish independent isogenized lines. To confirm deletions of the target genes, we performed genome DNA extraction and PCR amplification among these 50 lines by using primers listed in Table S4. Out of 50 lines, several lines from each target pair possessed deletion mutations. The PCR products were sequenced, and complete deletions were confirmed in all lines. We selected one mutant allele for each target gene and named them as *EcI-2*^*1*^, *EcI-3*^*1*^, *EcI-4*^*1*^, and *EcI-3-4*^*1*^. *EcI-2*^*1*^ has a 3704-bp deletion including the 5’ untranslated region and almost the entire *EcI-2/Oatp33Ea* CDS (Figure S2A). *EcI-3*^*1*^ has a 3535-bp deletion including the 5’ untranslated region and almost the entire *EcI-3/Oatp58Dc* CDS (Figure S2B). *EcI-4*^*1*^ has two deletion with a 9-bp deletion including the transcriptional start site and a 2092-bp deletion including almost the entire *EcI-4/Oatp58Db* CDS (Figure S2C). *EcI-3-4*^*1*^ has a 6495-bp deletion including the 5’ untranslated region of *EcI-3/Oatp58Dc*, the entire *EcI-3/Oatp58Dc* CDS and almost the entire *EcI-4/Oatp58Db* CDS (Figure S2D).

### Pupariation timing analysis in *Drosophila*

Flies were allowed to lay eggs on 3% agar plates containing 30% grape juice (Welch) at 25°C. Newly hatched larvae were transferred into vials with mashed standard food (25 larvae/vial), and pupae were counted at indicated time points. For each genotype, 4 vials were scored 2 times a day.

### Pupal volume measurements in *Drosophila*

Pupal length and width were determined from images captured with a Zeiss Axiocam 506 color digital camera attached to a SteREO Discovery.V12 microscope (Zeiss). Images were processed using ImageJ 1.53v (NIH). The pupal volume was determined by the following approximate equation: 4/3π (length x width^2^).

### Egg laying assay in *Drosophila*

Four-day-old single virgin females and 5-6 males were transferred to a vial with standard food to allow mating and egg laying. After 24 hours, individual females were then transferred to a fresh vial for further egg laying for another 24 hours. The total number of eggs laid in 2 days (48 hours) was counted.

### Mutagenesis in *Aedes*

sgRNAs were designed immediately downstream of the predicted start codon of each ecdysone importer gene (Figure S4). One guanine was added at the 5′ terminal end of each sgRNA to facilitate transcription by T7 RNA polymerase (Table S5). Potential off-target binding was checked by using two online tools: https://zifit.partners.org/ZiFiT/ and https://crispr.mit.edu. dsDNA templates for sgRNA synthesis were generated by template-free PCR, using a specific forward primer for each gene and one universal reverse primer (Table S5). sgRNA was synthesized using the MEGAscript T7 Transcription Kit (Ambion) and purified using the MEGAclear Transcription Clean-Up Kit (Ambion) following the manufacturer’s protocols. A mixture of sgRNA (40 ng/μl) and Cas9 protein with NLS (PAN Bio; 300 ng/μl) was microinjected into the posterior pole of pre-blastoderm embryos at an angle of 10–25°. The sgRNA/Cas9 ratio was optimized according to a previously described protocol (27). The embryos were hatched 5 days post injection and reared thereafter as described above. The microinjection was repeated three times for a total of 450 embryos per each gene.

The genomic DNA was isolated from newly hatched larvae using a buffer containing 1 M Tris-HCL, 0.5 M EDTA, 5 M NaCl, and Proteinase K (20 mg/ml). Fragments containing sgRNA target sites were amplified using gene-specific primers (Table S5). The PCR products were gel-purified using Gel DNA Recovery Kit (Zymoclean), cloned into the pGEMT-easy vector (Thermo Fisher Scientific), and sequenced. CRISPR/Cas9 mutagenesis efficiency was assessed through a T7 Endonuclease 1 (T7E1) assay. In brief, 200 ng of PCR products in 1X NEB buffer 2 (New England Biolabs) were hybridized under the following conditions: 95°C for 5 min, 95-85°C at −2°C/s, 85-25°C at −1°C/s, and held at 4°C. Ten units of T7E1 enzyme (New England Biolabs) were added to each sample, and the samples were incubated at 37°C for 15 min. Products were visualized using 2% agarose gel electrophoresis.

### Double-strand RNA (dsRNA) synthesis for RNAi in *Aedes*

For dsRNA synthesis, a T7 promoter sequence, TAATACGACTCACTATAGGGAGA, was added to the 5’ end of each primer as listed in Table S5. PCR was performed using the OneTaq-Quick-Load 2X Master Mix (New England BioLabs) with *Aedes* whole-body cDNA as a template, and the amplified PCR products were cleaned using the Gel DNA Recovery Kit (Zymoclean). dsRNA was synthesized using the MEGAscript T7 Transcription Kit (Ambion).

### RNAi in *Aedes* larvae

Prior to hatching, mosquito eggs were allowed to develop for a minimum of one week. To collect newly hatched larvae, eggs were submerged in a container with water and food and left at room temperature overnight. On the following day, the container was moved into the incubator set at 27°C and 80% relative humidity. 20 newly hatched larvae were transferred into 1.5 ml Eppendorf tube containing 100 μl of premixed food and dsRNA (0.5 μg/μl final concentration). Larvae were soaked in the dsRNA solution for 4 hours, after which they were transferred into a 12 oz cup containing 4 ml of premixed food. All surviving larvae were periodically monitored and counted thereafter. The head capsule length (head size) and siphon length were measured using images captured with a Zeiss Axiocam 506 color digital camera attached to a SteREO Discovery.V12 microscope (Zeiss).

For 20E and CF rescue experiments, 20E (Sigma-Aldrich; final concentration of 1 μM in 2% ethanol), CF (Sigma-Aldrich; final concentration of 10 nM in 2% ethanol), or 2% ethanol (control) was added into each cup 22 hours after hatching, and surviving larvae were periodically monitored and counted thereafter.

### RNAi in *Aedes* adult females

Purified dsRNA was resuspended with HPLC-grade water (Thermo Fisher Scientific) at 5 μg/μl. Newly emerged virgin females were separated using manual aspirator into a new cage and provided with 10% sucrose. Three days after eclosion, virgin females were anesthetized on ice and injected with 2.0 μg dsRNA (400 nl) using a Nanoject II microinjector (Drummond Scientific Company). After injection, females were placed back in a cage with 10% sucrose to recover for 24 hours at 27°C with 80% humidity.

### Ovary length, follicle size, and yolk length measurements in *Aedes* adult females

After post-injection recovery, adult females were allowed to mate for 48 hours and then allowed to feed on bovine blood for 1 hour. Fully engorged females were selected, and isolated in a cage and provided with 10% sucrose. At 24 hours post blood meal, dsRNA-injected females were anesthetized for 5 min at −20°C. Ovaries were then dissected, and the length of the central longitudinal axis of the ovary was measured. The ovary length of each female was calculated as an average length of the two ovaries. For primary follicle size measurement, the length of the longitudinal axis of fifteen primarily follicles per ovary was measured, and the average follicle length of each female was calculated. The yolk length of each female was determined by measuring the average yolk length of all oocytes from one ovary. All measurements were conducted using images captured with a Zeiss Axiocam 506 color digital camera attached to a SteREO Discovery.V12 microscope (Zeiss). Images were processed using ImageJ 1.53v (NIH).

### Egg laying assay in *Aedes*

After post-injection recovery, adult females were allowed to mate for 48 hours and then allowed to feed on bovine blood for 1 hour. Fully engorged females were selected, and individual females were placed in a cup with oviposition paper. The number of eggs was counted at 96 hours post blood meal.

## ACKNOWLEDGEMENTS

We thank A.S. Raikhel for his generous support in establishing our mosquito research project; Bloomington *Drosophila* Stock Center (NIH P40 OD018537) and T. Nishimura for fly stocks; M.E. Adams for cell lines; *Drosophila* Genomics Resource Center (NIH P40 OD010949), National Institute of Genetics Fly Stock Center, and D.J. Mangelsdorf for vectors and cDNA clones; and A.S. Raikhel and M.E. Adams for critical reading of the manuscript. This study was supported by a Postdoctoral Fellowship for Research Abroad from the Japan Society for the Promotion of Science to N.O., the Naito Foundation Subsidy for Dispatch of Young Researchers Abroad to N.O., an NIH Director’s New Innovator Award DP2 GM132929 to N.Y., a research grant from the W.M. Keck Foundation to N.Y., and a Pew Biomedical Scholars Award from the Pew Charitable Trusts to N.Y.

## Supplementary Information

**Figure S1.**
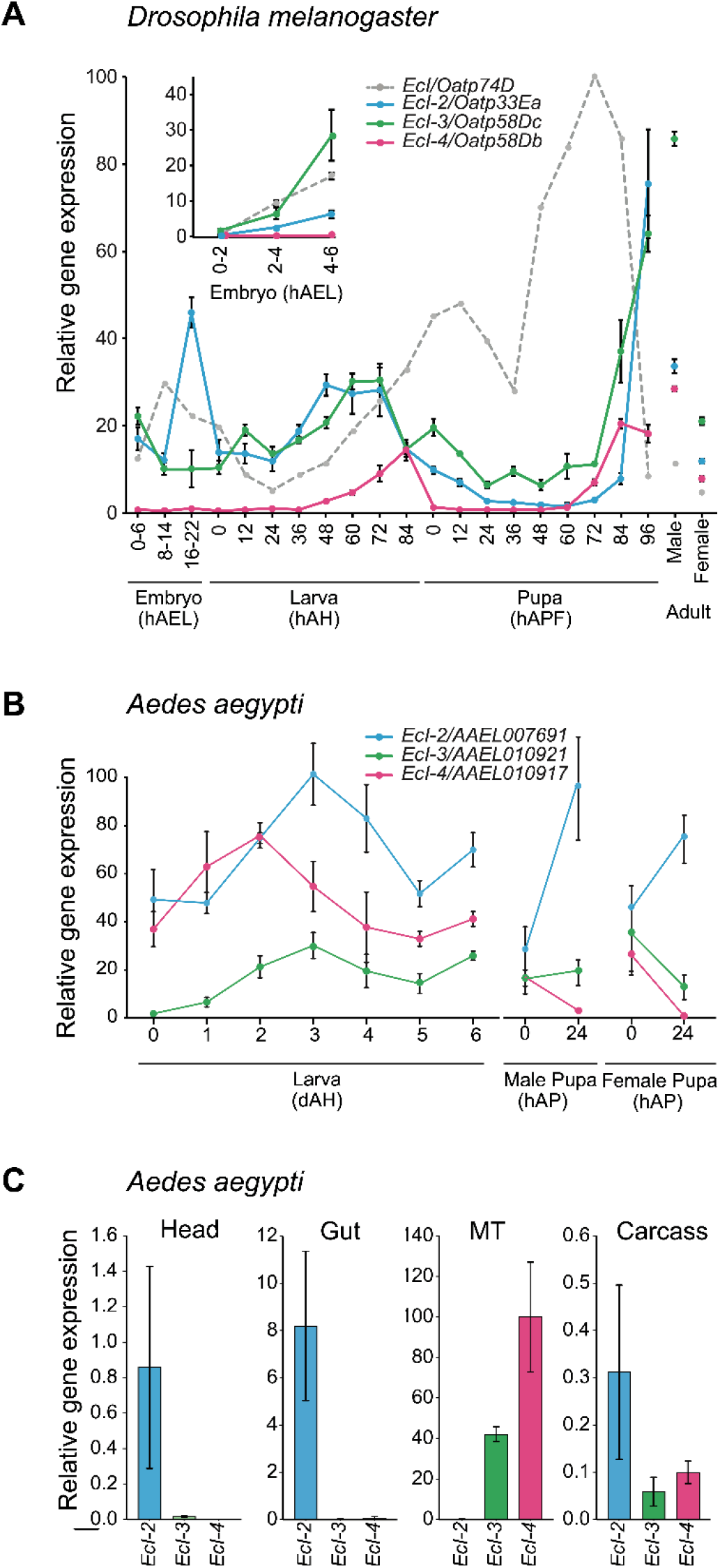
Expression levels of ecdysone importer genes in *Drosophila* and *Aedes*. (A) Relative expression levels of ecdysone importer genes in the whole body during *Drosophila* development, as assessed by qRT-PCR. Samples were collected from *w1118* animals. *EcI* expression levels are adopted from Okamoto et al. 2018, and values are shown as percentages relative to the highest expression level of *EcI*. hAEL, hours after egg laying, hAH, hours after hatching, hAPF, hours after puparium formation. Adult cDNA samples were prepared from flies at 24 hr after eclosion. All values are the means ± SD (n = 3). (B) Relative expression levels of ecdysone importer genes in the whole body during *Aedes* development, as assessed by qRT-PCR. Values are shown as percentages relative to the highest expression level of *EcI-2*. dAH, days after hatching, hAP, hours after pupation. All values are the means ± SEM (n = 3). (C) Relative expression levels of ecdysone importer genes in various tissues in *Aedes*, as assessed by qRT-PCR. Tissues were dissected from 4th instar larvae. Values are shown as percentages relative to the expression level of *EcI-4* in the Malpighian tubule. MT, Malpighian tubule. All values are the means ± SEM (n = 3).

**Figure S2.**
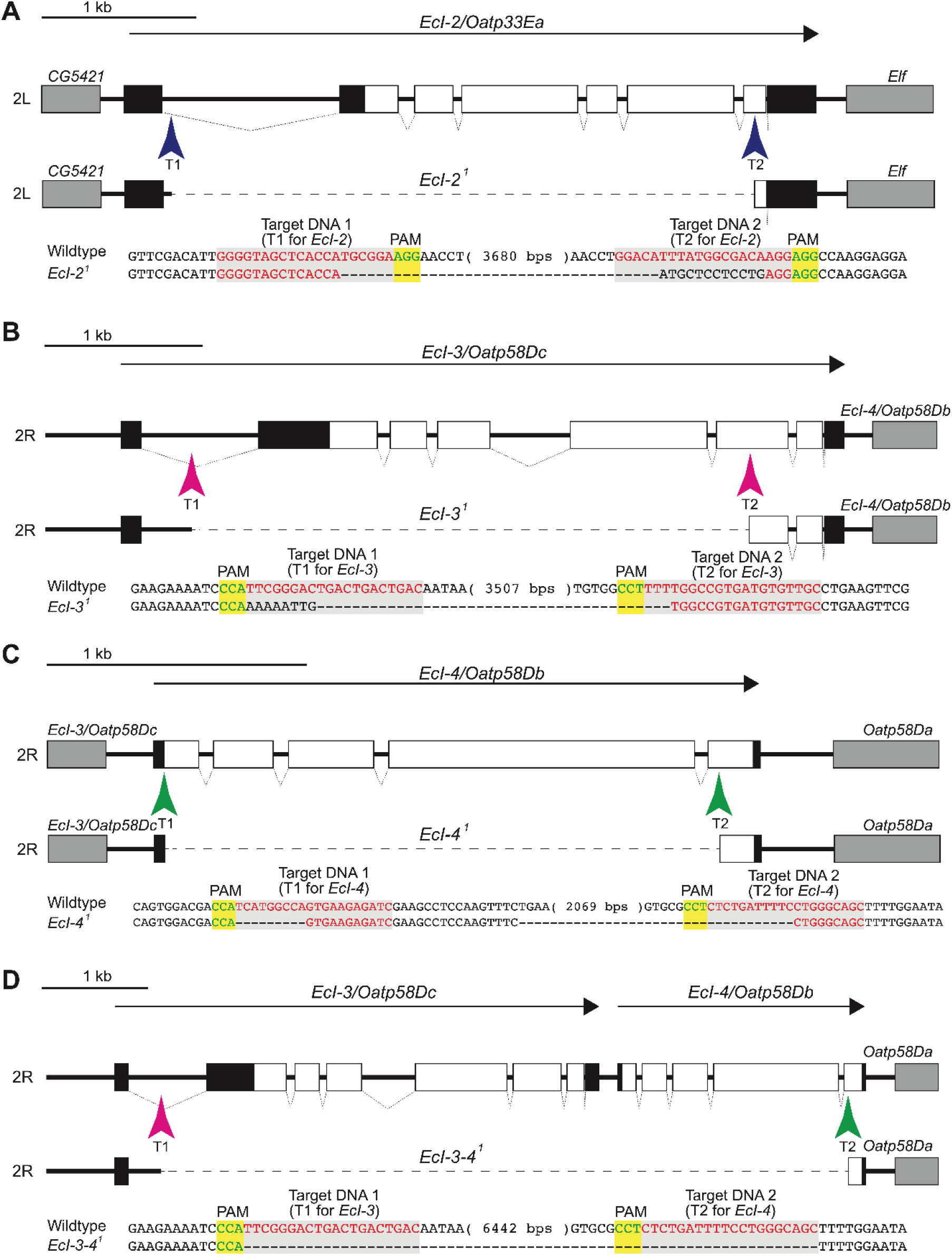
Generation of ecdysone importer mutants in *Drosophila* using CRISPR/Cas9. (A-D) Schematic representation of the guide RNA (gRNA) targets for generating *EcI-2* (A), *EcI-3* (B), *EcI-4* (C), and *EcI-3-4* double (D) mutants in *Drosophila*. The protein-coding DNA sequences (CDSs) and untranslated regions are represented by open and filled boxes, respectively. Neighboring genes are represented by gray boxes. Arrows indicate the orientation of ecdysone importer genes, while arrowheads (T1 and T2) indicate gRNA target sequences. Sequences of ecdysone importer mutants as compared to the wildtype sequences are shown at the bottom of each panel. gRNA target sequences are shown in red, and the neighboring NGG protospacer adjacent motif (PAM) sequences are shown in green. Deleted residues are shown as dashes.

**Figure S3.**
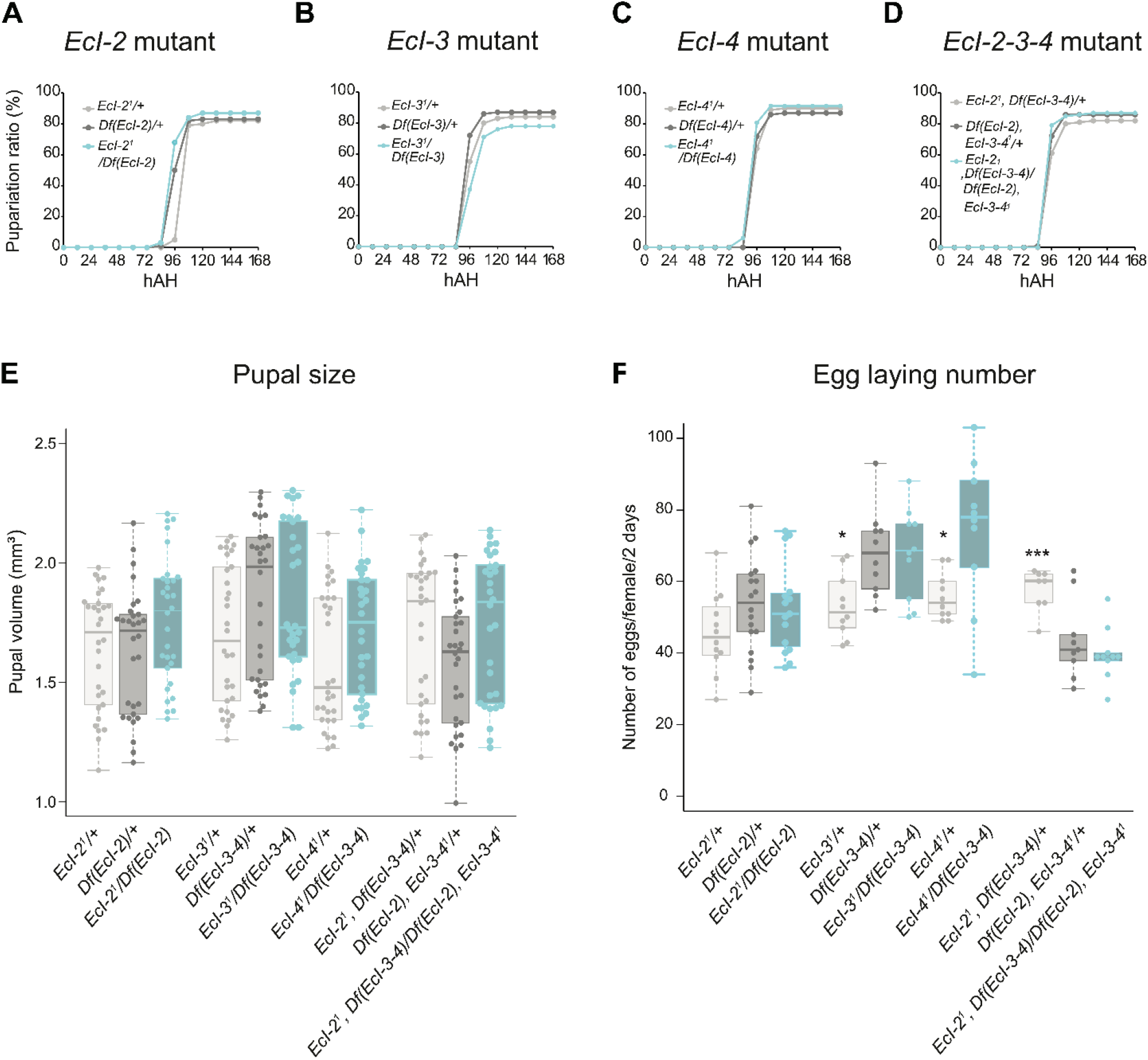
Growth and reproductive phenotypes of ecdysone importer mutants in *Drosophila*. (A-D) Pupariation timing of *EcI-2* (A), *EcI-3* (B), *EcI-4* (C), and *EcI-2-3-4* triple (D) mutants in *Drosophila*. Df indicates deficiency alleles over ecdysone importer genes indicated in parenthesis. Blue lines indicate transheterozygous ecdysone importer mutants, whereas gray lines indicate heterozygous controls. hAH, hours after hatching. n = 100 from 4 independent experiments. (E, F) Box plot of pupal size (E) and egg laying number (F) of transheterozygous ecdysone importer mutants (blue) and heterozygous controls (gray). Pupal size is shown as pupal volume (mm3) calculated using the length and width of the pupae (n = 30 from 4 independent experiments), whereas egg laying number is shown as the number of eggs laid per female every two days (n = 10-18 from 2 independent experiments). *p < 0.05, ***p < 0.001 from Mann-Whitney U test with Bonferroni Correction.

**Figure S4.**
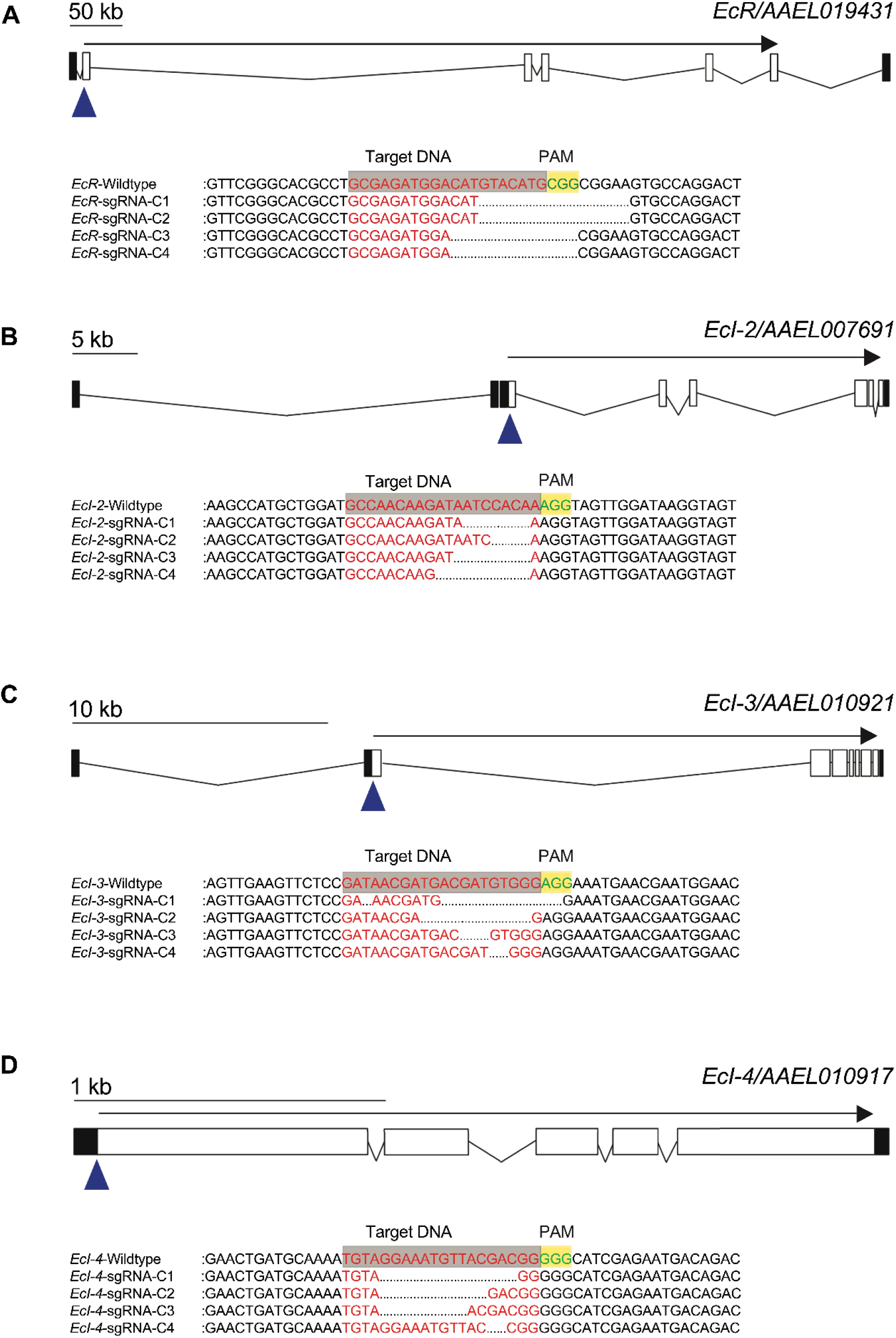
Generation of *EcR* and ecdysone importer mutants in *Aedes* using CRISPR/Cas9. (A-D) Schematic representation of the guide RNA (gRNA) targets for generating *EcR* (A), *EcI-2* (B), *EcI-3* (C), and *EcI-4* (D) mutants in *Aedes*. The protein-coding DNA sequences (CDSs) and untranslated regions are represented by open and filled boxes, respectively. Arrows indicate the orientation of the genes, while arrowheads indicate gRNA target sequences. Sequences of the mutants from 4 independent PCR-amplified clones are shown as compared to the wildtype sequences at the bottom of each panel. gRNA target sequences are shown in red, and the neighboring NGG protospacer adjacent motif (PAM) sequences are shown in green. Deleted residues are shown as dotted lines.

**Figure S5.**
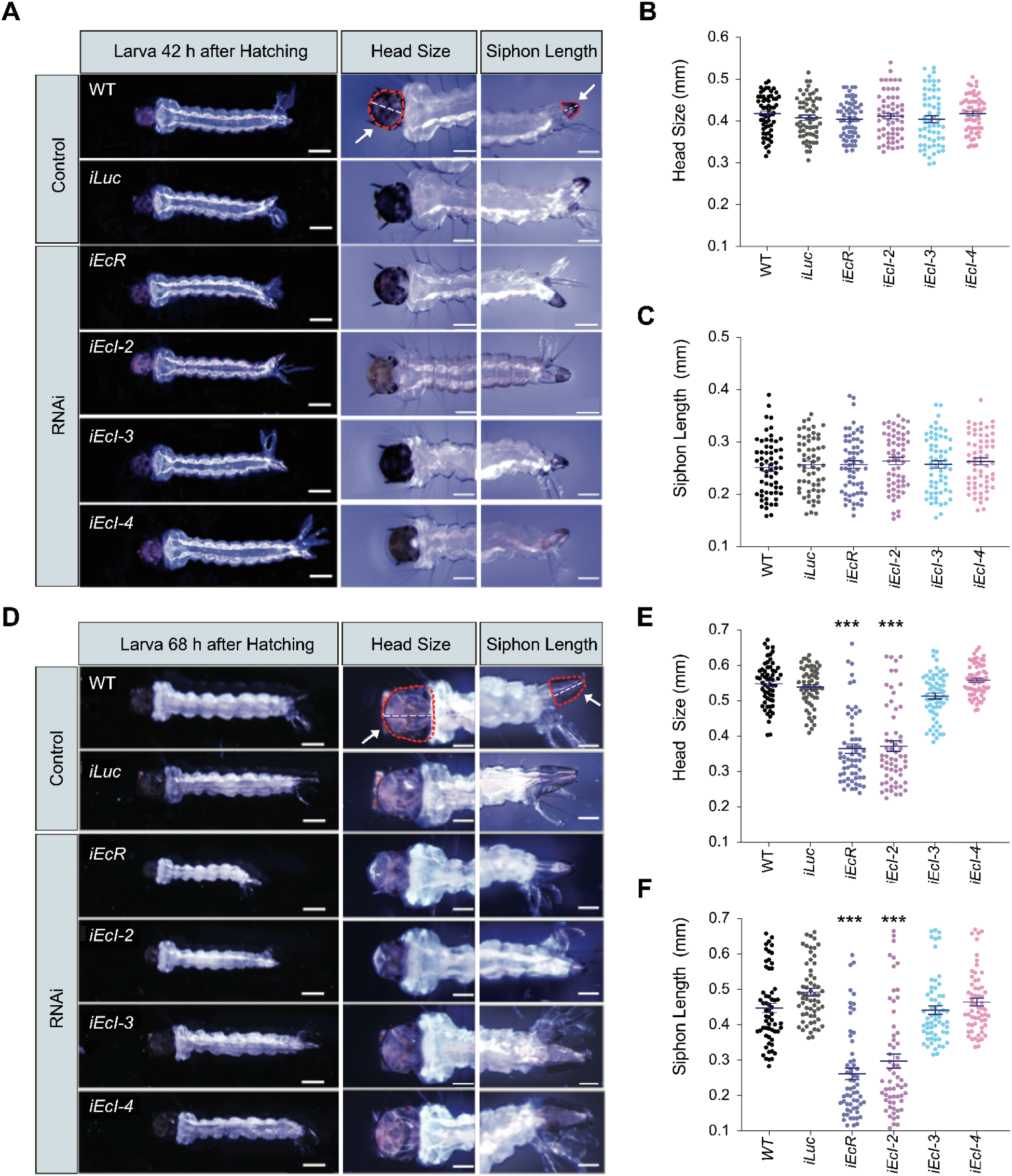
Detailed developmental phenotype of dsRNA-treated *Aedes* larvae. (A-F) Representative images of the whole body, head, and siphon of dsRNA-treated larvae and their measurements at 42 hrs (A-C) or 68 hrs (D-F) after hatching. WT, wildtype (control); *iLuc*, *Luc* RNAi (control); *iEcR*, *EcR* RNAi; *iEcI-2*, *EcI-2* RNAi; *iEcI-3*, *EcI-3* RNAi; *iEcI-4*, *EcI-4* RNAi. Scale bars, 0.5 mm for whole larva images; 250 μm for head and siphon images. The head and siphon are circled with red dashed lines and marked with white arrows in the WT images in (A) and (D). The head capsule length (head size) and siphon length (indicated by white dashed lines in the same images) were measured in (B), (C), (E), and (F). All values are the means ± SEM (n = 60 from 3 independent experiments). ***p < 0.001 from one-way ANOVA followed by Dunnett’s multiple comparison test as compared to *iLuc* control.

**Figure S6.**
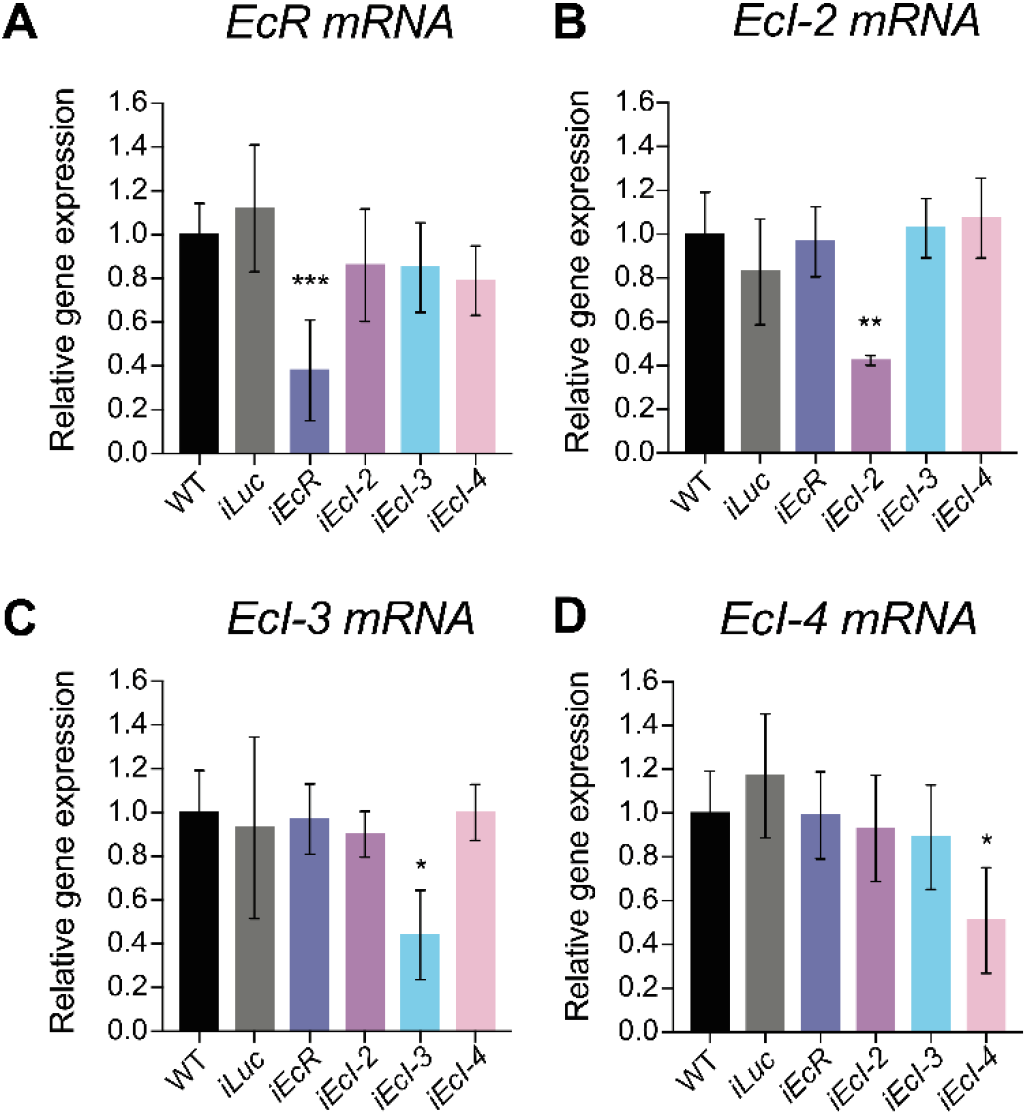
Knockdown efficiency and specificity of dsRNA soaking experiments in *Aedes* larvae. (A-D) Relative expression levels of *EcR* (A), *EcI-2* (B), *EcI-3* (C), and *EcI-4* (D) in WT (wildtype; control), *Luc* RNAi (*iLuc*; control), *EcR* RNAi (*iEcR*), and *EcI-2*, *3*, and *4* RNAi (*iEcI-2*, *iEcI-3*, and *iEcI-4*) animals at 48 hours after hatching, as assessed by qRT-PCR. All values are the means ± SEM (n = 4). *p < 0.05, **p < 0.01, ***p < 0.001 from one-way ANOVA followed by Dunnett’s multiple comparison test as compared to *iLuc* control.

**Figure S7.**
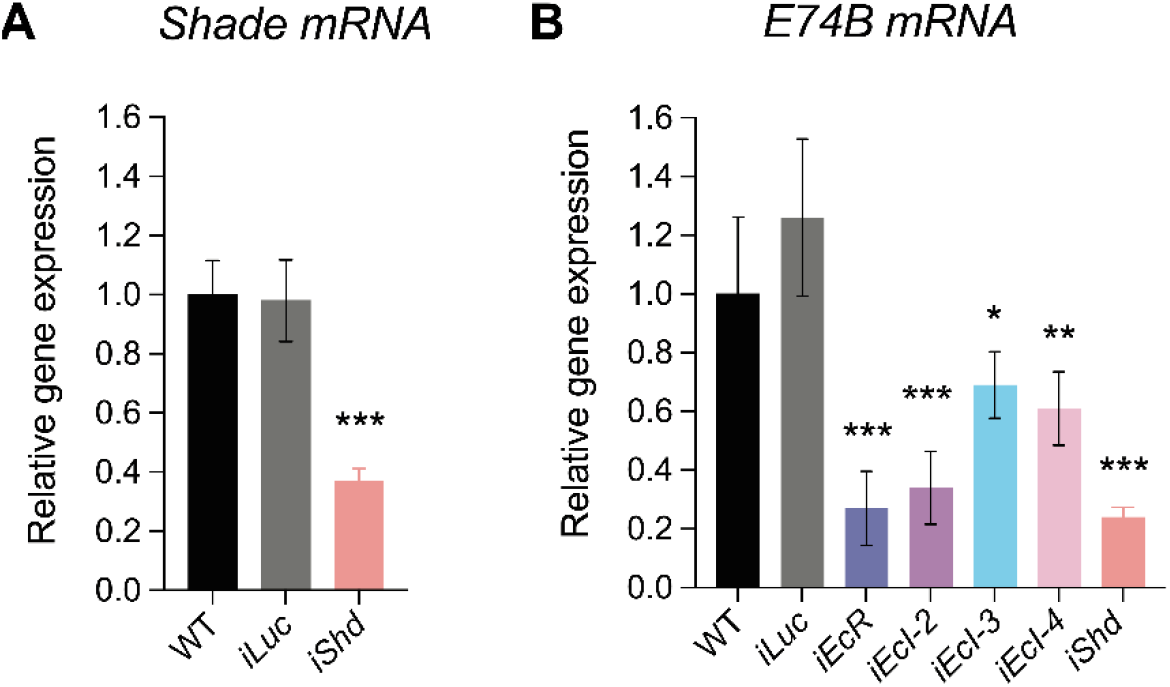
*Shade* and *E74B* expression levels in dsRNA-treated *Aedes* larvae. (A) Relative expression levels of *Shade* in WT (wildtype; control), *Luc* RNAi (*iLuc*; control), and *Shade* RNAi (*iShd*) animals at 48 hours after hatching, as assessed by qRT-PCR. All values are the means ± SEM (n = 3). ***p < 0.001 from one-way ANOVA followed by Dunnett’s multiple comparison test as compared to *iLuc* control. (B) Relative expression levels of *E74B* in WT (wildtype; control), *Luc* RNAi (*iLuc*; control), *EcR* RNAi (*iEcR*), *EcI-2*, *3*, and *4* RNAi (*iEcI-2*, *iEcI-3*, and *iEcI-4*), and *Shade* RNAi (*iShd*) animals at 48 hrs after hatching, as assessed by qRT-PCR. All values are the means ± SEM (n = 4). *p < 0.05, **p < 0.01, ***p < 0.001 from one-way ANOVA followed by Dunnett’s multiple comparison test as compared to *iLuc* control.

**Figure S8.**
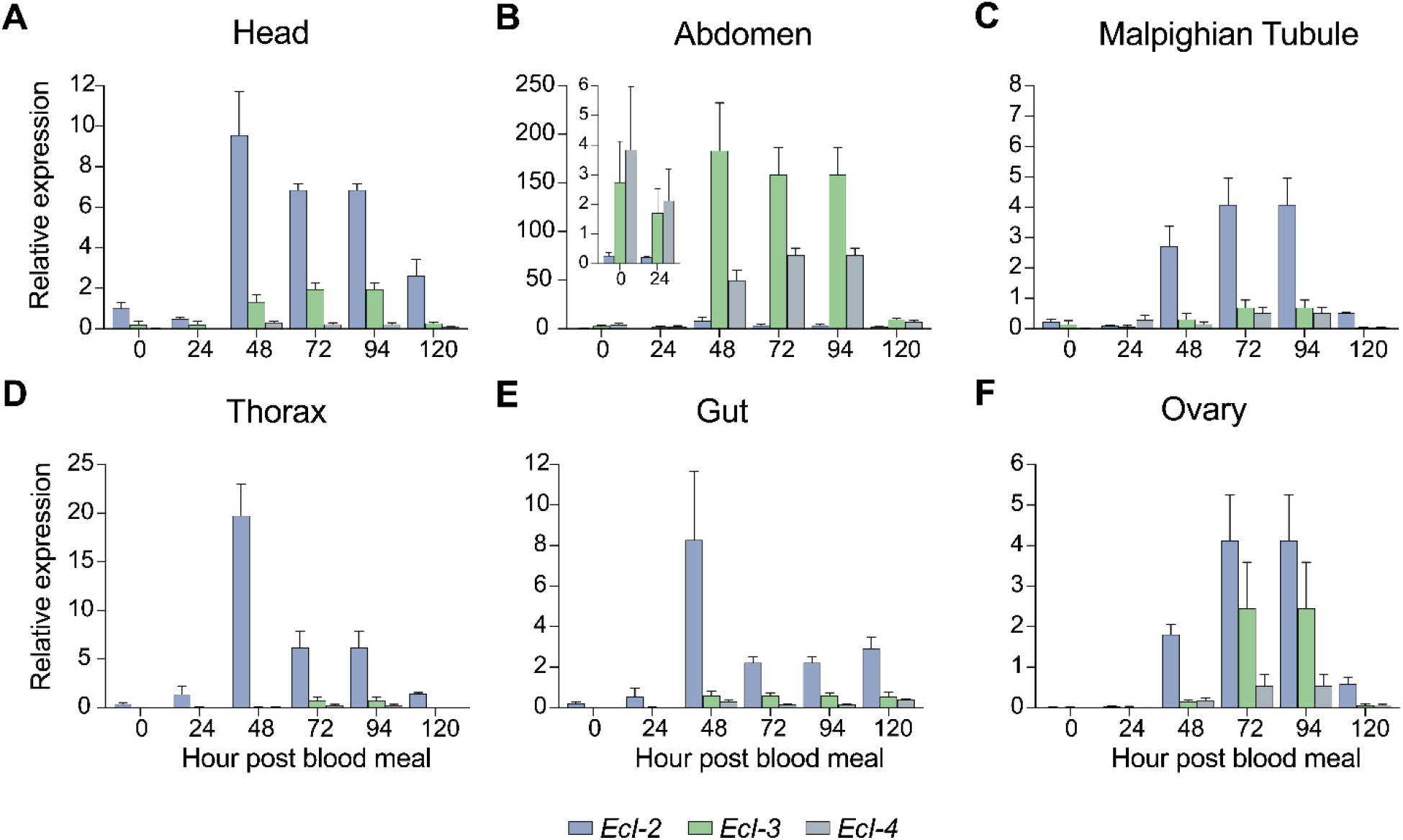
Ecdysone importer gene expression in *Aedes* adult females after blood meal. (A-F) Relative expression levels of *EcI-2*, *3*, and *4* in the head (A), abdomen (B), Malpighian tubule (C), thorax (D), gut (E), and ovary (F), as assessed by qRT-PCR. Values are shown relative to the *EcI-2* level in the head. All values are the means ± SEM (n = 5).

**Figure S9.**
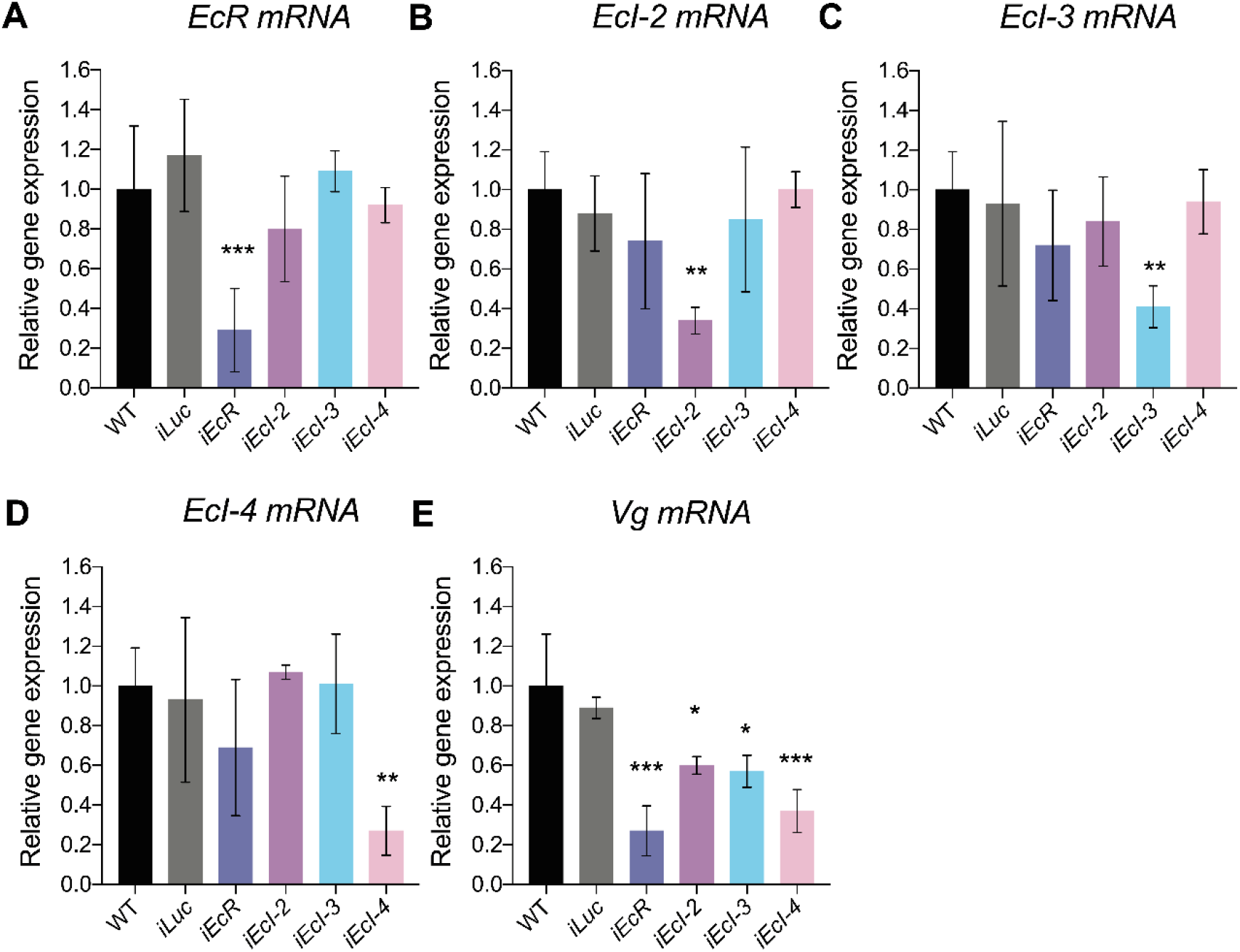
Gene expression levels in dsRNA-injected *Aedes* adult females. (A-E) Relative expression levels of *EcR* (A), *EcI-2* (B), *EcI-3* (C), *EcI-4* (D), and *Vg* (E) in the fat body (abdominal wall with adhered fat body) of WT (wildtype; control), *Luc* RNAi (*iLuc*; control), *EcR* RNAi (*iEcR*), and *EcI-2*, *3*, and *4* RNAi (*iEcI-2*, *iEcI-3*, and *iEcI-4*) adult females at 24 hours post blood meal, as assessed by qRT-PCR. All values are the means ± SEM (n = 3). *p < 0.05, **p < 0.01, ***p < 0.001 from one-way ANOVA followed by Dunnett’s multiple comparison test as compared to *iLuc* control.

**Table S1.** Embryonic lethality in *Drosophila* ecdysone importer mutants.

| Cross # | Genes Mutated |  |  |  | Expected Homozygous Larval Ratio (%) | Number of Larvae Analyzed | Homozygous Larval Number | Homozygous Larval Ratio (%) | Embryonic Hatchability (%) |
| --- | --- | --- | --- | --- | --- | --- | --- | --- | --- |
|  | *Ecl* | *Ecl-2* | *Ecl-3* | *Ecl-4* |  |  |  |  |  |
| 1 | - | + | + | + | 33.33 | 217 | 72 | 33.18 | 99.5 |
| 2 | + | - | + | + | 50.00 | 184 | 91 | 49.46 | 98.9 |
| 3 | + | + | - | + | 50.00 | 182 | 93 | 51.10 | 102.2 |
| 4 | + | + | + | - | 50.00 | 178 | 93 | 52.25 | 104.5 |
| 5 | + | - | - | + | 33.33 | 228 | 73 | 32.02 | 96.1 |
| 6 | + | - | + | - | 33.33 | 117 | 43 | 36.75 | 110.3 |
| 7 | + | + | - | - | 50.00 | 176 | 88 | 50.00 | 100.0 |
| 8 | + | - | - | - | 33.33 | 139 | 46 | 33.09 | 99.3 |
| 9 | - | - | + | + | 16.67 | 409 | 75 | 18.34 | 110.0 |
| 10 | - | + | - | + | 16.67 | 286 | 47 | 16.43 | 98.6 |
| 11 | - | + | + | - | 16.67 | 154 | 28 | 18.18 | 109.1 |
| 12 | - | - | - | + | 11.11 | 290 | 30 | 10.34 | 93.1 |
| 13 | - | - | + | - | 11.11 | 124 | 13 | 10.48 | 94.4 |
| 14 | - | + | - | - | 16.67 | 193 | 33 | 17.10 | 102.6 |
| 15 | - | - | - | - | 11.11 | 122 | 13 | 10.66 | 95.9 |

**Table S2.**
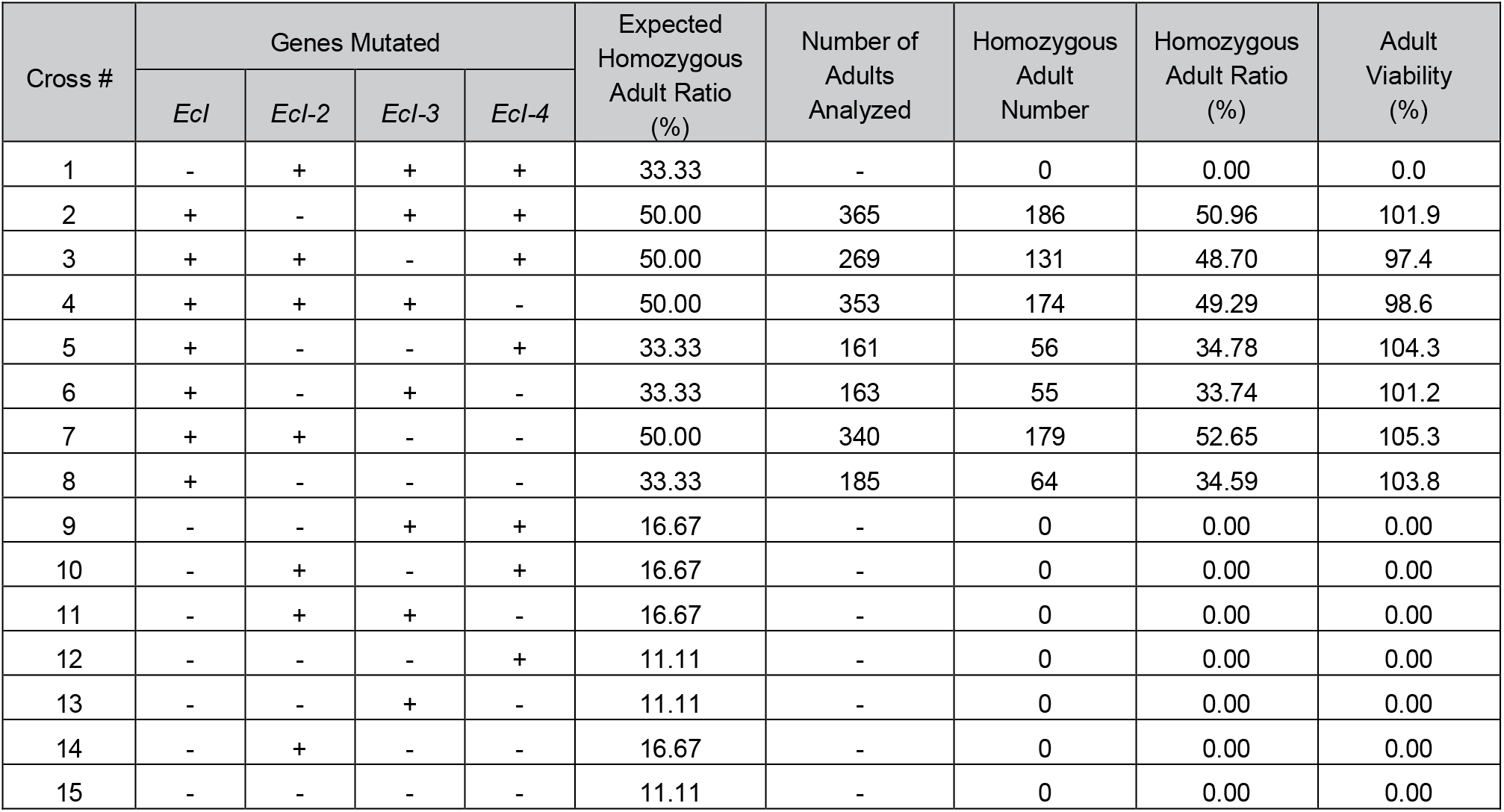
Post-embryonic lethality in *Drosophila* ecdysone importer mutants.

**Table S3.** OATP proteins used for phylogenetic analysis.

| Species | Protein name / Gene ID | GenBank accession number |
| --- | --- | --- |
| *Drosophila melanogaster* | Ecl/Oatp74D | NP_648989 |
|  | Ecl-2/Oatp33Ea | NP_609568 |
|  | Ecl-3/Oatp58Dc | NP_611659 |
|  | Ecl-4/Oatp58Db | NP_611658 |
|  | Oatp26F | NP_609055 |
|  | Oatp30B | NP_723463 |
|  | Oatp33Eb | NP_609570 |
|  | Oatp58Da | NP_611657 |
| *Musca domestica* | MDOA001191 | XP_005183411 |
|  | MDOA012869 | XP_011295428 |
|  | MDOA000381 | XP_005176681 |
|  | MDOA000437 | XP_005176680 |
|  | MDOA011793 | XP_005176679 |
|  | MDOA005021 | XP_011292793 |
|  | MDOA012671 | XP_005184218 |
|  | MDOA001872 | XP_005174920 |
| *Phlebotomus papatasi* | PPAI001606 | N.A. |
|  | PPAI000868 (Pseudogene) | N.A. |
|  | PPAI000869 | N.A. |
|  | PPAI006402 | N.A. |
|  | PPAI006000 | N.A. |
|  | PPAI004257 | N.A. |
|  | PPAI005206 (Pseudogene) | N.A. |
|  | PPAI001455 | N.A. |
| *Aedes aegypti* | Ecl-2/AAEL007691 | XP_001658583 |
|  | Ecl-3/AAEL010921 | XP_001661189 |
|  | Ecl-4/AAEL010917 | XP_001661188 |
|  | AAEL001812 | XP_001660406 |
|  | AAEL001808 | XP_001660407 |
|  | AAEL007693 | XP_001658582 |
| *Anopheles gambiae* | AGAP010042 | XP_319187 |
|  | AGAP006638 | XP_316669 |
|  | AGAP006637 | XP_316668 |
|  | AGAP008712 | XP_314819 |
|  | AGAP008711 | XP_557860 |
|  | AGAP010043 | XP_319188 |
| *Culex quinquefasciatus* | CPIJ006275 | XP_001847741 |
|  | CPIJ005186 | XP_001847079 |
|  | CPIJ005183 | XP_001847076 |
|  | CPIJ001972 | XP_001843749 |
|  | CPIJ001971 | XP_001843748 |
|  | CPIJ001970 | XP_001843747 |
|  | CPIJ006274 | XP_001847740 |

**Table S4.**
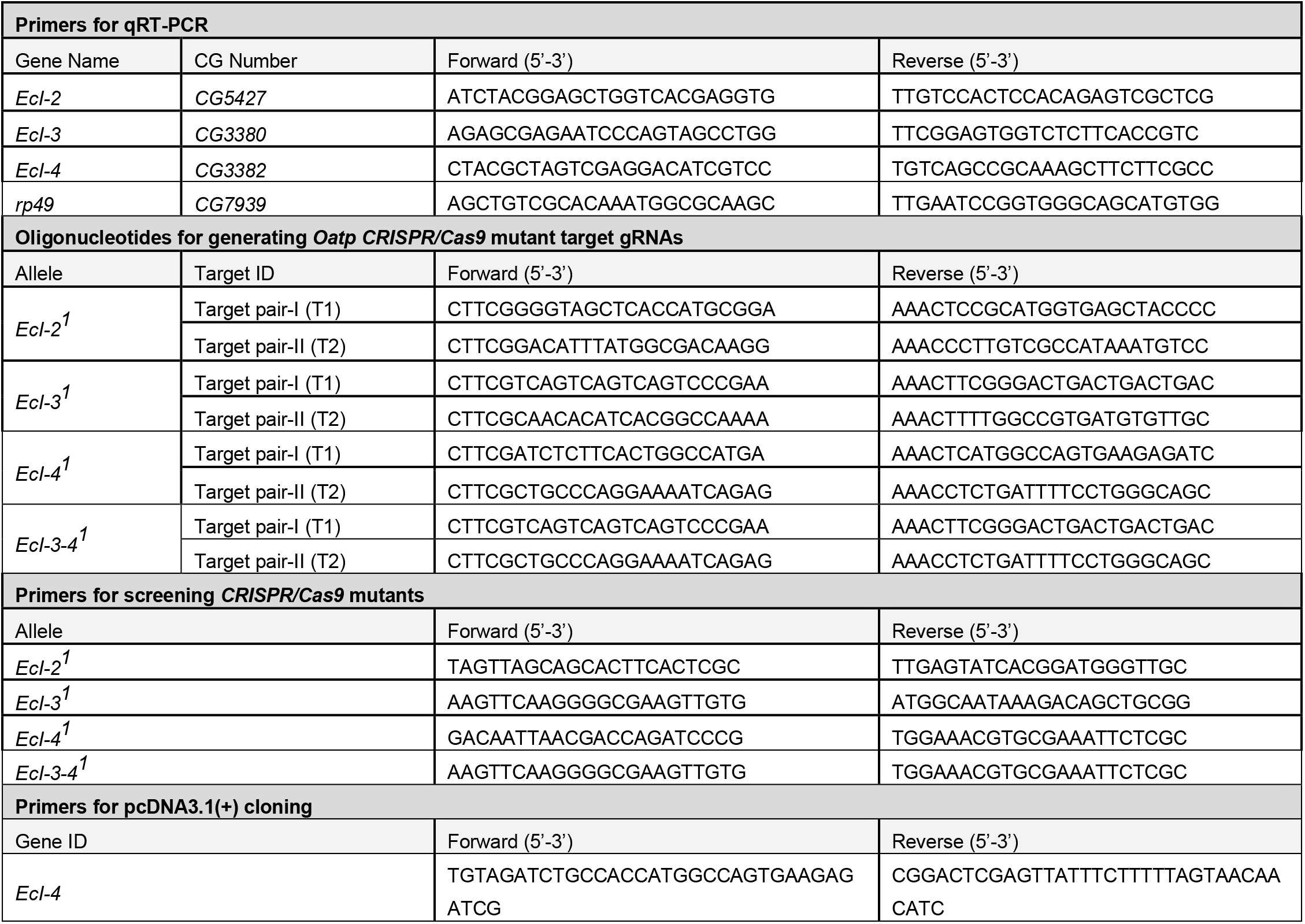
Oligonucleotides used for *Drosophila* study.

**Table S5.**
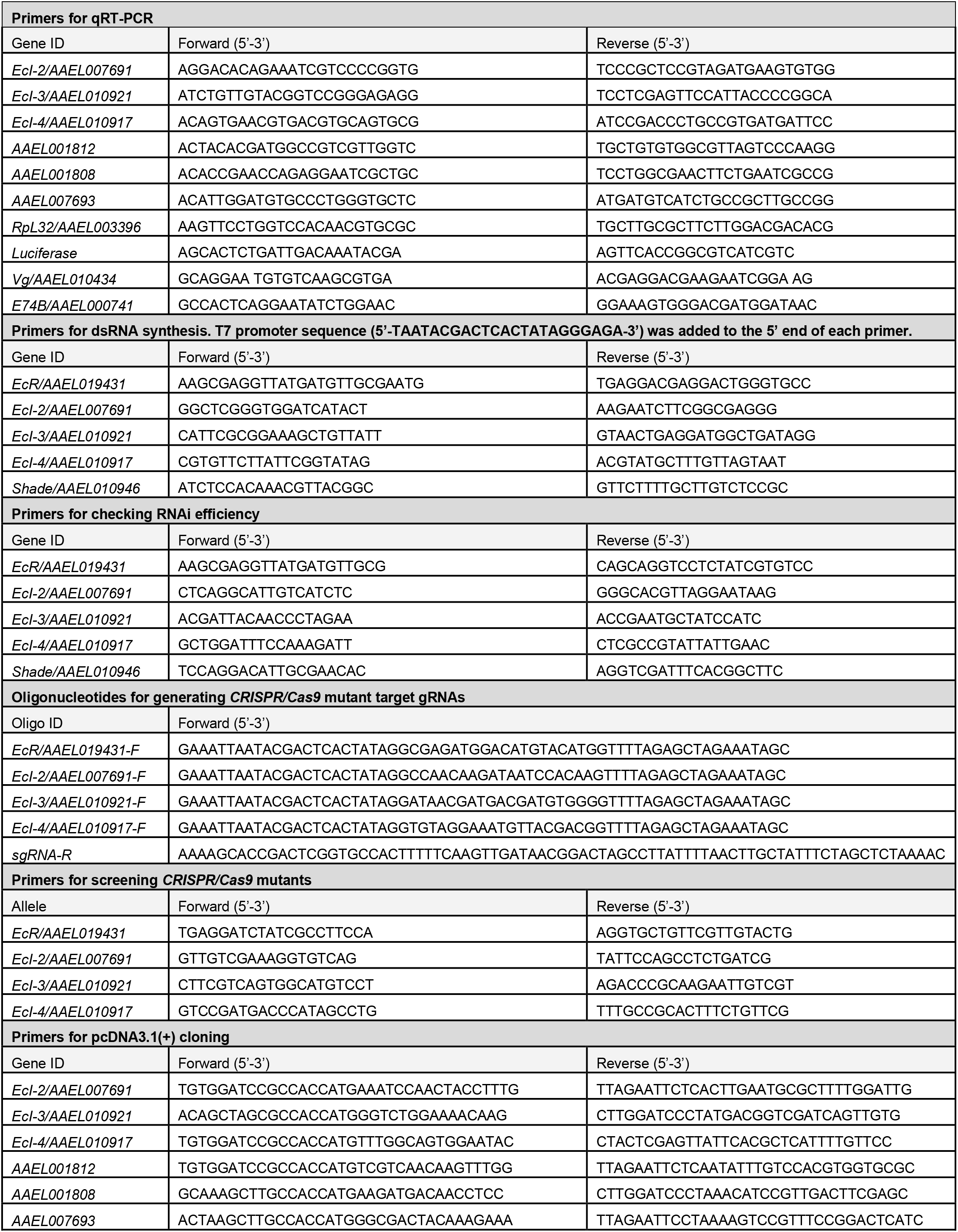
Oligonucleotides used for *Aedes* study.

## REFERENCES

1. N. Yamanaka, Ecdysteroid signalling in insects—From biosynthesis to gene expression regulation. Adv Insect Physiol 60, 1–36 (2021).

2. H. E. Thomas, H. G. Stunnenberg, A. F. Stewart, Heterodimerization of the *Drosophila* ecdysone receptor with retinoid X receptor and ultraspiracle. Nature 362, 471–475 (1993).

3. T.-P. Yao, W. A. Segraves, A. E. Oro, M. McKeown, R. M. Evans, *Drosophila* ultraspiracle modulates ecdysone receptor function via heterodimer formation. Cell 71, 63–72 (1992).

4. T.-P. Yao, et al., Functional ecdysone receptor is the product of EcR and Ultraspiracle genes. Nature 366, 476–479 (1993).

5. L. M. Riddiford, P. Cherbas, J. W. Truman, Ecdysone receptors and their biological actions. Vitam Horm 60, 1–73 (2000).

6. N. Okamoto, et al., A membrane transporter is required for steroid hormone uptake in *Drosophila*. Dev Cell 47, 294–305.e7 (2018).

7. J. Rösner, J. Tietmeyer, H. Merzendorfer, Organic anion-transporting polypeptides are involved in the elimination of insecticides from the red flour beetle, *Tribolium castaneum*. J Pest Sci, (2021).

8. M. U. Kraemer, et al., The global distribution of the arbovirus vectors *Aedes aegypti* and *Ae. albopictus*. Elife 4, e08347 (2015).

9. S. Leta, et al., Global risk mapping for major diseases transmitted by *Aedes aegypti* and Aedes albopictus. Int J Infect Dis 67, 25–35 (2018).

10. S. Roy, T. T. Saha, Z. Zou, A. S. Raikhel, Regulatory pathways controlling insect reproduction. Annu Rev Entomol 63, 1–23 (2017).

11. I. A. Hansen, G. M. Attardo, S. D. Rodriguez, L. L. Drake, Four-way regulation of mosquito yolk protein precursor genes by juvenile hormone-, ecdysone-, nutrient-, and insulin-like peptide signaling pathways. Front Physiol 5, 103 (2014).

12. L. Chen, J. Zhu, G. Sun, A. S. Raikhel, The early gene Broad is involved in the ecdysteroid hierarchy governing vitellogenesis of the mosquito *Aedes aegypti*. J Mol Endocrinol 33, 743–761 (2004).

13. A. S. Raikhel, Vitellogenesis in mosquitoes. Adv Dis Vector Res, 1–39 (1992).

14. B. J. Matthews, et al., Improved reference genome of *Aedes aegypti* informs arbovirus vector control. Nature 563, 501–507 (2018).

15. A. Domingos, R. Pinheiro‐Silva, J. Couto, V. do Rosário, J. de la Fuente, The *Anopheles gambiae* transcriptome – a turning point for malaria control. Insect Mol Biol 26, 140–151 (2017).

16. W. F. S. Martins, et al., Transcriptomic analysis of insecticide resistance in the lymphatic filariasis vector *Culex quinquefasciatus*. Sci Rep 9, 11406 (2019).

17. D. E. Neafsey, et al., Mosquito genomics. Highly evolvable malaria vectors: the genomes of 16 *Anopheles* mosquitoes. Science 347, 1258522 (2015).

18. M. V. Sharakhova, et al., Update of the *Anopheles gambiae* PEST genome assembly. Genome Biol 8, R5 (2007).

19. P. Arensburger, et al., Sequencing of *Culex quinquefasciatus* establishes a platform for mosquito comparative genomics. Science 330, 86–88 (2010).

20. K. S. Christopherson, M. R. Mark, V. Bajaj, P. J. Godowski, Ecdysteroid-dependent regulation of genes in mammalian cells by a *Drosophila* ecdysone receptor and chimeric transactivators. Proc Natl Acad Sci USA 89, 6314–6318 (1992).

21. D. No, T. P. Yao, R. M. Evans, Ecdysone-inducible gene expression in mammalian cells and transgenic mice. Proc Natl Acad Sci USA 93, 3346–3351 (1996).

22. E. Saez, et al., Identification of ligands and coligands for the ecdysone-regulated gene switch. Proc Natl Acad Sci USA 97, 14512–14517 (2000).

23. A. Petryk, et al., Shade is the *Drosophila* P450 enzyme that mediates the hydroxylation of ecdysone to the steroid insect molting hormone 20-hydroxyecdysone. Proc Natl Acad Sci USA 100, 13773–13778 (2003).

24. K. F. Rewitz, L. I. Gilbert, Daphnia Halloween genes that encode cytochrome P450s mediating the synthesis of the arthropod molting hormone: Evolutionary implications. BMC Evol Biol 8, 60 (2008).

25. N. Okamoto, N. Yamanaka, Steroid hormone entry into the brain requires a membrane transporter in *Drosophila*. Curr Biol 30, 359–366.e3 (2020).

26. S. Kondo, R. Ueda, Highly improved gene targeting by germline-specific Cas9 expression in *Drosophila*. Genetics 195, 715–721 (2013).

27. K. E. Kistler, L. B. Vosshall, B. J. Matthews, Genome engineering with CRISPR-Cas9 in the mosquito *Aedes aegypti*. Cell Rep 11, 51–60 (2015).

28. S. R. Christophers, *Aedes aegypti* (L.), the yellow fever mosquito. Its life history, bionomics, and structure. Cambridge Univ. Press, Cambridge (1960).

29. A. Bar. and J. Andrew, Morphology and morphometry of *Aedes aegypti* larvae. Annu Res Rev Biol 3, 1–12 (2013).

